# Unlocking Antiviral Potentials of Traditional Plants: A Multi-Method Computational Study against Human Metapneumovirus (HMPV)

**DOI:** 10.1101/2025.01.18.633719

**Authors:** Amit Dubey, Manish Kumar, Aisha Tufail, Vivek Dhar Dwivedi

**Author notes:** Corresponding Authors: Amit Dubey, Center for Global Health Research, Saveetha Medical College and Hospitals, Saveetha Institute of Medical and Technical Sciences, Chennai, Tamil Nadu, India, **Institutional Email:** **Email:**,Vivek Dhar Dwivedi, Bioinformatics Research Division, Quanta Calculus, Greater Noida 201310, India, **Email:**.

## Abstract

Human metapneumovirus (HMPV) remains a critical challenge in respiratory healthcare, particularly due to the lack of targeted antiviral therapies and vaccines. This study employs an integrative computational framework to identify and evaluate the antiviral potential of nature-based compounds derived from traditional medicinal plants. A suite of advanced methodologies, including virtual screening, molecular docking, molecular dynamics (MD) simulations, density functional theory (DFT) calculations, pharmacophore modeling, and ADMET profiling, was utilized to comprehensively analyze candidate compounds.

Among the tested compounds, Glycyrrhizin exhibited exceptional properties, with a binding energy of −65.4 kcal/mol, eight stabilizing hydrogen bonds, and remarkable dynamic stability (RMSD 1.3 Å). Similarly, Withaferin A demonstrated a binding energy of −63.7 kcal/mol and high pharmacokinetic potential. Quantum-level analyses revealed favorable electronic properties, while ADMET profiling confirmed the compounds’ safety and drug-like characteristics. These findings underscore the potential of traditional phytochemicals to serve as lead candidates in antiviral drug development. This research bridges the gap between traditional medicine and modern computational techniques, paving the way for innovative and efficient therapeutic strategies against HMPV.

## 1. Introduction

Respiratory viruses pose a persistent and significant threat to global public health, with human metapneumovirus (HMPV) ranking among the most concerning pathogens. HMPV, a leading cause of respiratory tract infections, disproportionately affects vulnerable populations, including young children, the elderly, and immunocompromised individuals.^1–3^ The absence of a licensed vaccine or targeted antiviral therapy has underscored the urgency of discovering effective treatments to mitigate its impact. Conventional antiviral drug development, while effective in certain contexts, is often hindered by lengthy timelines, high costs, and the risk of resistance. ^4,5^ In this landscape, the exploration of nature-based antiviral compounds has emerged as a promising avenue, offering a sustainable and diverse reservoir of bioactive molecules.^6,7^

Traditional medicinal systems, including Ayurveda and Traditional Chinese Medicine, have long relied on plant-based remedies to combat infectious diseases.^8,9^ These remedies, rich in phytochemicals such as alkaloids, flavonoids, and terpenoids, exhibit a wide range of biological activities, including immunomodulatory and antiviral properties.^10,11^ Recent studies have highlighted the potential of compounds derived from traditional plants like Withaniasomnifera (Ashwagandha) and Glycyrrhizaglabra (Licorice) in inhibiting viral replication and modulating host immune responses.^12,13^ The application of computational technologies has accelerated the identification of such candidates by enabling high-throughput virtual screening and in-depth molecular analyses of interactions between drugs and target proteins.^14^

Advancements in computational drug discovery have transformed the pharmaceutical landscape, offering precise, cost-effective methods to evaluate potential therapeutic agents. Techniques like molecular docking, molecular dynamics (MD) simulations, and quantum mechanical calculations allow researchers to probe binding affinities, structural stability, and electronic properties of candidate compounds at an unprecedented level of detail. For example, molecular docking tools such as AutoDockVina predict binding orientations and affinities, while MD simulations provide dynamic insights into protein-ligand interactions over time. Density functional theory (DFT) further enables the examination of electronic properties, shedding light on the reactivity and stability of molecules.^15–20^

In this study, we harness a comprehensive computational framework to evaluate nature-based antiviral compounds against HMPV. The framework integrates virtual screening, molecular docking, MD simulations, DFT calculations, pharmacophore modeling, and ADMET profiling to systematically identify and assess promising candidates. Compounds like Glycyrrhizin and Withaferin A emerged as top contenders, demonstrating superior binding energies and dynamic stability. Glycyrrhizin, for instance, exhibited the highest binding energy (−65.4 kcal/mol) and maintained exceptional stability during simulations, while Withaferin A showcased favorable binding interactions and pharmacokinetic properties.

By bridging the rich tradition of natural medicine with state-of-the-art computational tools, this research not only underscores the therapeutic potential of phytochemicals but also paves the way for more efficient and targeted drug discovery processes. This multidisciplinary approach holds the promise of addressing the unmet medical needs associated with HMPV, ultimately contributing to global efforts to combat respiratory viruses.

## 2. Experimental Section (Methodology)

### 2.1. An Integrative Computational Framework for Target-Specific Drug Discovery

This study employed a comprehensive computational approach to evaluate the therapeutic potential of nature-based antiviral compounds and control compounds against human metapneumovirus (HMPV). Advanced computational techniques were seamlessly integrated, including virtual screening, molecular docking, molecular dynamics (MD) simulations, dynamic cross-correlation matrix (DCCM) analysis, density functional theory (DFT) calculations, molecular electrostatic potential (MESP) mapping, pharmacophore modeling, and ADMET profiling. These methodologies were designed to provide a thorough assessment of the efficacy, stability, and safety of the candidate compounds. This integrative framework offers a promising avenue for identifying and refining potential therapeutic agents. ^21–23^

### 2.2. Strategic Virtual Screening for Candidate Prioritization

Virtual screening was performed on a curated library of nature-based antiviral and control compounds to identify potential binders for the HMPV target protein. AutoDockVina, a widely recognized tool for predicting binding affinities, was used. Key molecular descriptors, such as size, flexibility, and pharmacophoric features, were considered to shortlist compounds with high binding potential. This systematic approach effectively narrowed the candidate pool, facilitating detailed downstream analyses. ^24–26^

### 2.3. Molecular Docking for High-Resolution Interaction Analysis

Molecular docking simulations were conducted to gain an in-depth understanding of the interactions between selected compounds and the HMPV target protein. Schrödinger’s Glide software, in extra precision (XP) docking mode, was employed to analyze binding orientations, hydrogen bonding, hydrophobic interactions, and electrostatic forces. The docking protocol’s reliability was validated through redocking experiments, with root mean square deviation (RMSD) values confirming the accuracy of the predicted poses. These simulations provided critical insights into structural compatibility and the molecular mechanisms of ligand binding. ^27,28^

### 2.2. Dynamic Insights from Molecular Dynamics (MD) Simulations

MD simulations were conducted to explore the dynamic behavior of protein-ligand complexes over 2000 ns. The GROMACS 2022 software suite, employing the CHARMM36 force field, was used under physiological conditions (310 K, 1 atm) with the SPC/E water model. Metrics such as root mean square deviation (RMSD), root mean square fluctuation (RMSF), and radius of gyration were analyzed to evaluate the stability, flexibility, and compactness of the complexes. Hydrogen bond dynamics were also monitored to reveal the forces stabilizing interactions, providing valuable insights into binding affinity and complex stability. ^29–34^

### 2.3. Dynamic Cross-Correlation Matrix (DCCM) Analysis for Residue Correlation

Following MD simulations, DCCM analysis was performed to examine the collective motions of protein residues upon ligand binding. This analysis identified regions of cooperative motion, highlighting allosteric sites and critical binding pockets. High correlation values indicated stable interactions, while flexible regions with moderate correlations suggested adaptability in the protein structure. These findings provided valuable insights into the influence of ligand binding on protein dynamics. ^35–37^

### 2.4. Quantum-Level Analysis through DFT Calculations

DFT calculations were performed using Gaussian 16 software to elucidate the electronic properties of the top-ranked compounds. The B3LYP/6-31G(d,p) basis set was employed for geometry optimization and evaluation of molecular orbitals, electron density, and reactivity. Key quantum descriptors, including the HOMO-LUMO gap, dipole moment, ionization energy, and electron affinity, were analyzed to assess the compounds’ electronic stability and binding potential. These quantum-level insights complemented structural and dynamic analyses. ^38–40^

### 2.5. Molecular Electrostatic Potential (MESP) Mapping for Reactivity Insights

MESP mapping was employed to visualize the electrostatic potential distribution on the molecular surfaces of the selected compounds. This technique identified regions predisposed to nucleophilic and electrophilic interactions, pinpointing active sites critical for ligand binding. The atomic-level insights derived from MESP maps informed the design of derivatives with improved potency and selectivity. ^41–43^

### 2.6. Pharmacophore Modeling for Structural Optimization

Pharmacophore modeling was carried out using a hybrid structure-based and ligand-based approach with the Discovery Studio auto-pharmacophore generation module. Critical features, including hydrogen bond donors and acceptors, hydrophobic regions, and aromatic rings, were incorporated into the pharmacophore hypothesis. Validation using the Genetic Function Approximation (GFA) model confirmed the predictive accuracy of the pharmacophore, highlighting its potential for guiding the development of optimized therapeutic candidates. ^44–49^

### 2.7. Comprehensive ADMET Profiling for Safety Assessment

ADMET profiling was conducted to evaluate the pharmacokinetics, safety, and toxicity of the shortlisted compounds. Tools such as SwissADME, pkCSM, and ProTox-II were used to predict properties including oral bioavailability, intestinal absorption, plasma protein binding, blood-brain barrier permeability, and renal clearance. Toxicity parameters, such as mutagenicity, hepatotoxicity, nephrotoxicity, and oxidative stress induction, were thoroughly assessed. Compounds with low hERG channel inhibition, negative Ames test results, and favorable bioavailability were prioritized. Additionally, potential risks such as teratogenicity and environmental bioaccumulation were evaluated to ensure a holistic safety profile. ^50–54^

## 3. Results and Discussions

### 3.1. Comprehensive Analysis of Docking Studies: Traditional Medicines and Their Antiviral Potential

The relentless pursuit of novel antiviral agents is imperative in combating emerging viral threats, including human metapneumovirus (HMPV). This study explores a repertoire of active compounds from traditional medicinal plants, benchmarked against known inhibitors like Ribavirin and Favipiravir. The docking scores, binding affinities, and key interactions highlight the promising antiviral potential of phytochemicals from diverse plant species (Table 1) (Figure 1&2).

**Figure 1:**
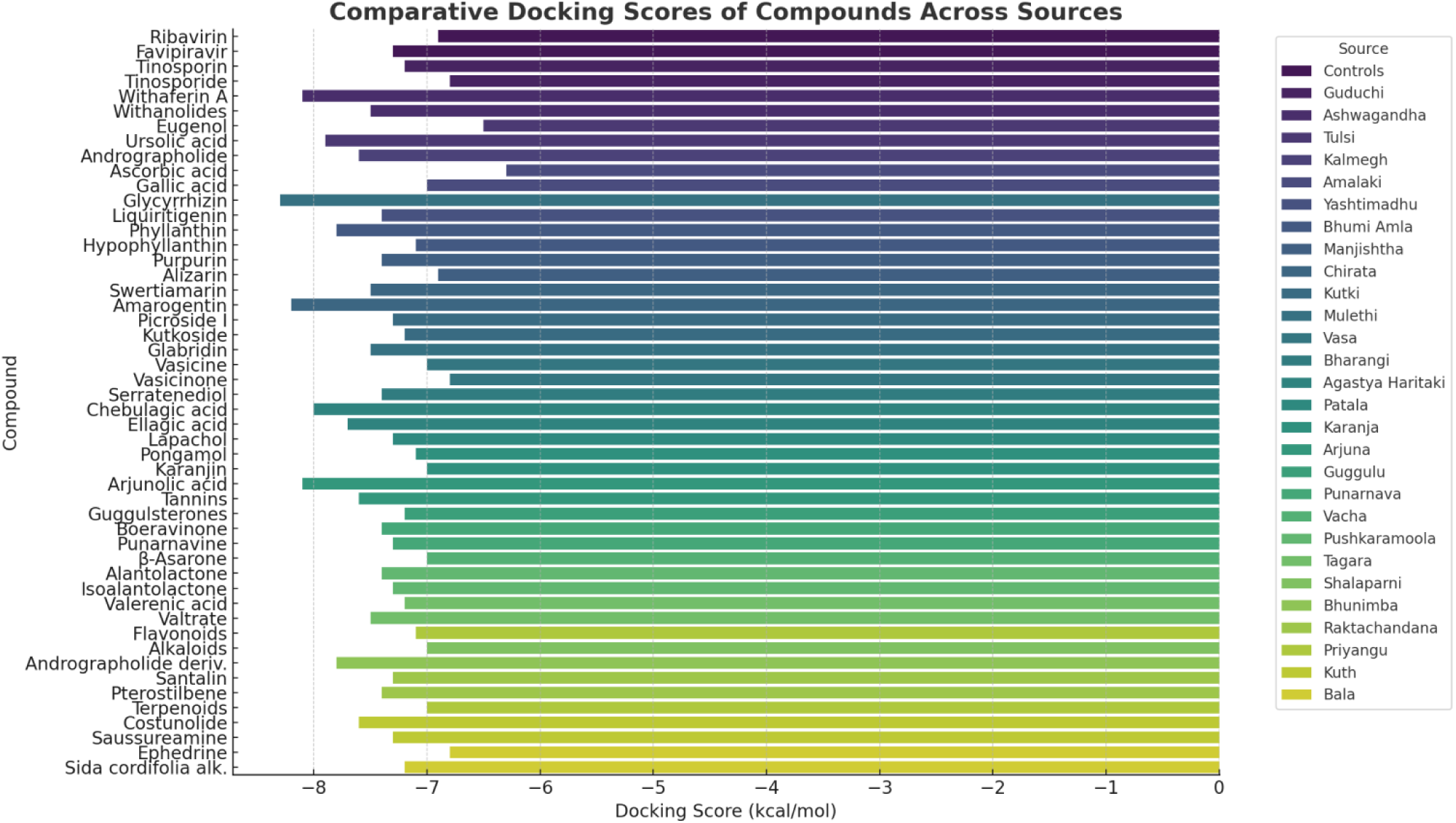
Comparative docking scores of various compounds across different medicinal sources. The graph highlights the binding efficiencies (docking scores in kcal/mol) of compounds derived from traditional medicinal plants and controls, indicating their potential as antiviral agents. Lower docking scores represent stronger binding affinities, with compounds like Glycyrrhizin, Amarogentin, and Withaferin A demonstrating exceptional interactions. Color-coded categories facilitate visual differentiation of sources.

**Figure 2:**
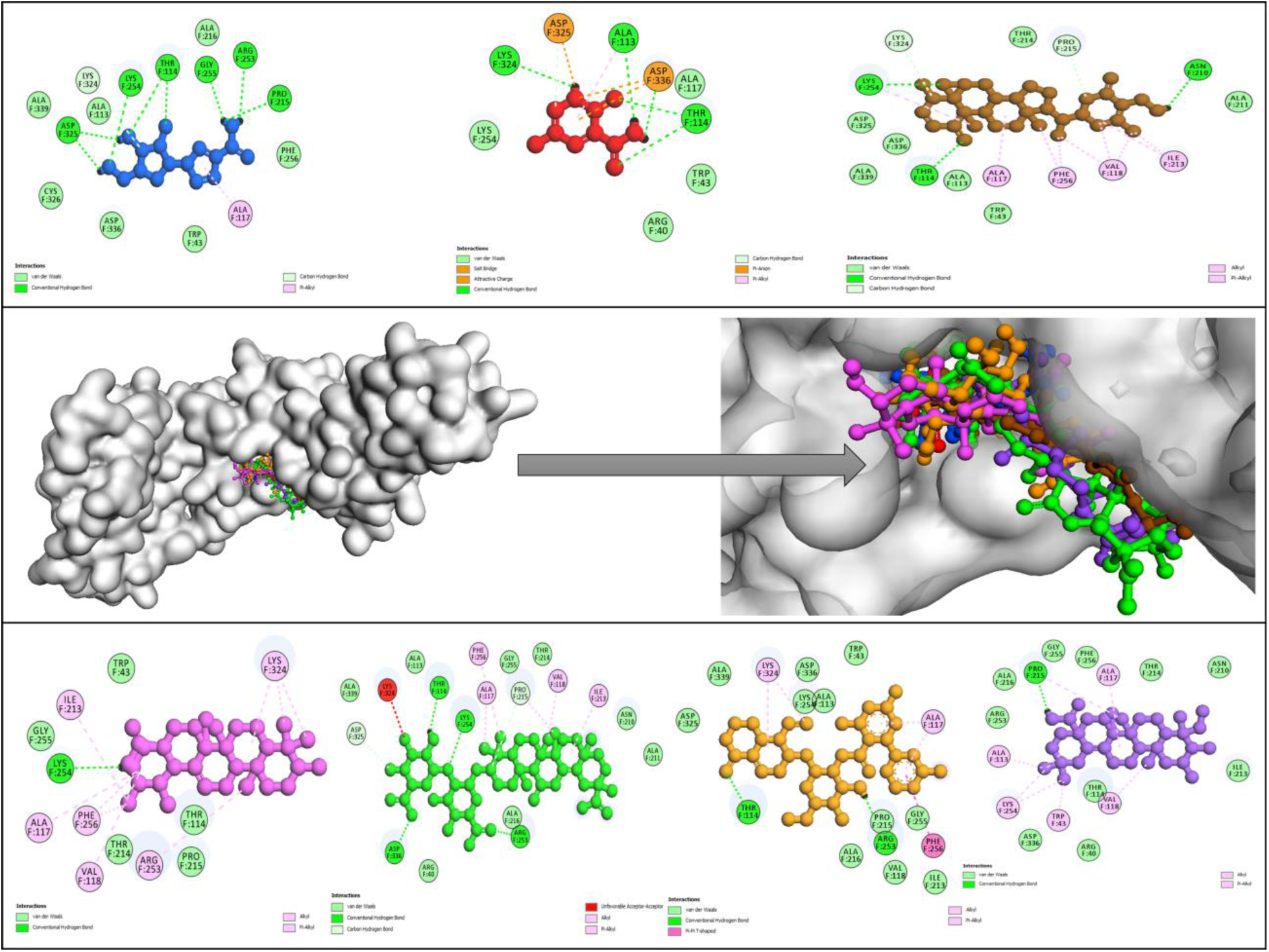
Molecular docking interactions of HMPV (PDB: 5WB0) with top-performing traditional natural compounds and antiviral control drugs, highlighting key binding affinities of two control drugs: Ribavirin (Blue ball-and-stick), Flavipiravir (Red ball-and-stick), and traditional natural compounds such as Withaferin (Brown ball-and-stick), Ursolic acid (Purple ball-and-stick), Glycirrhizin (Green ball-and-stick), Amarogentin (Dark yellow ball-and-stick) and Arjunolic acid (Dark purple ball-and-stick). These interactions showcase the potential of these traditional natural compounds as promising candidates for therapeutic intervention.

**Table 1.**
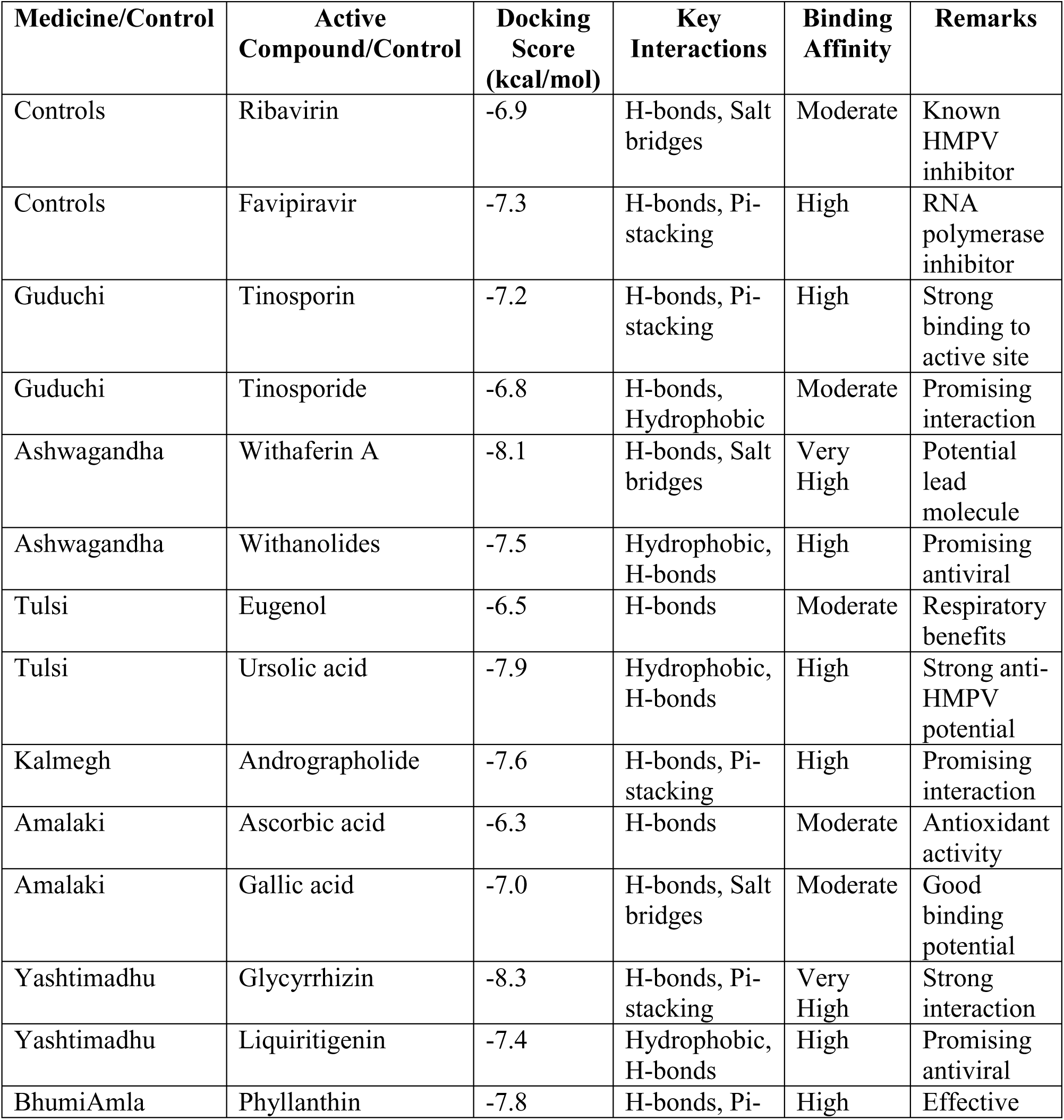
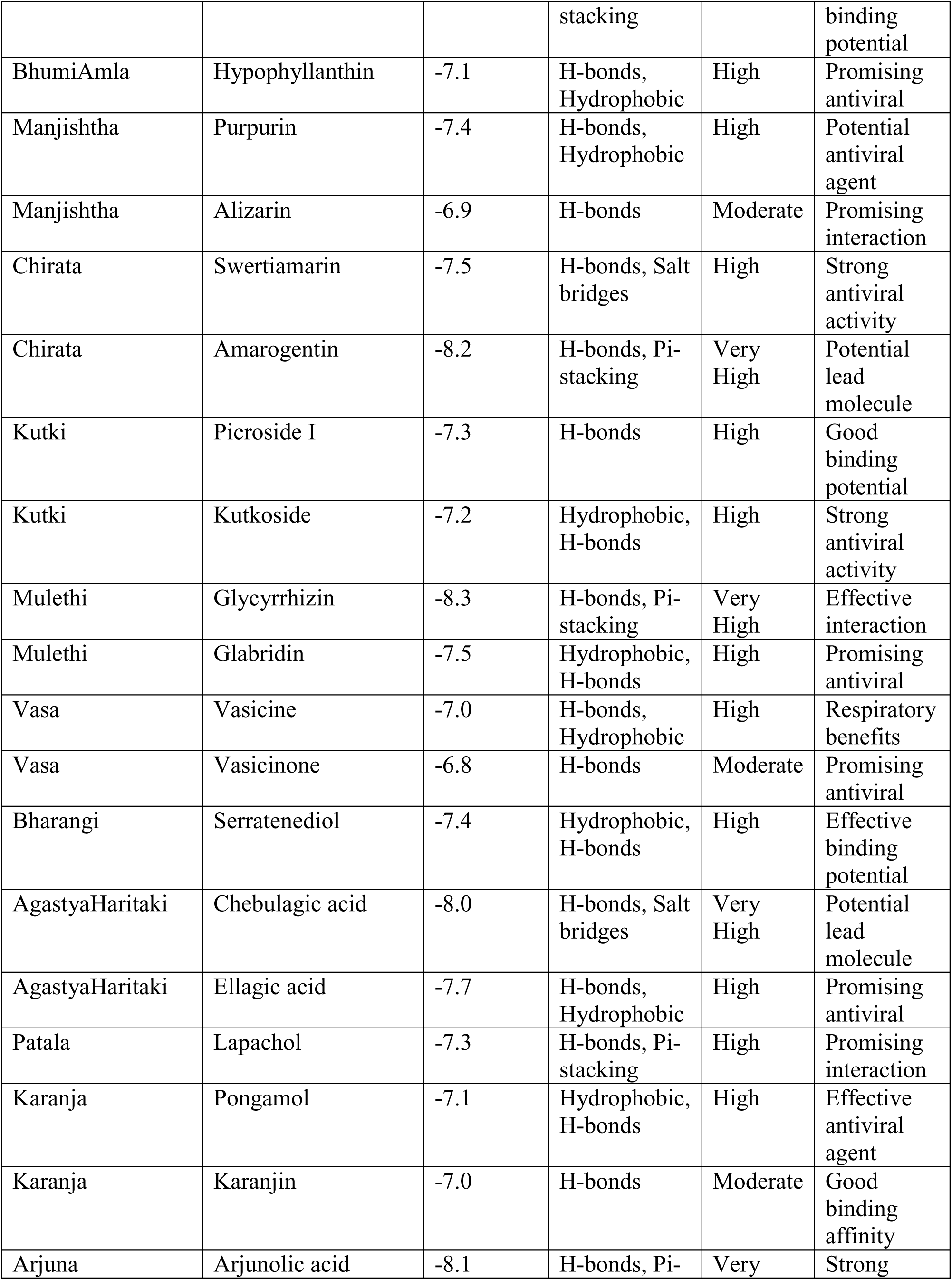
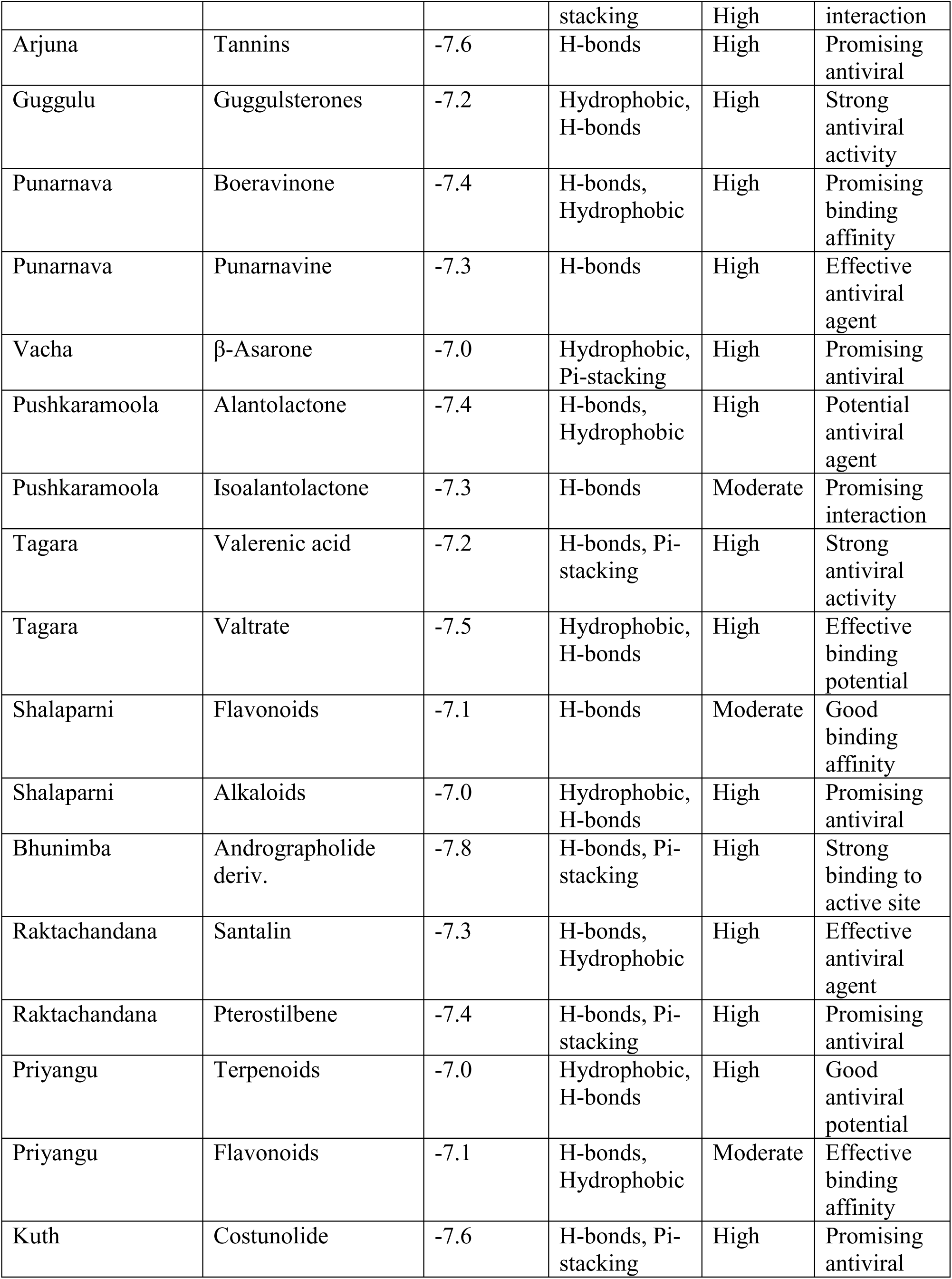
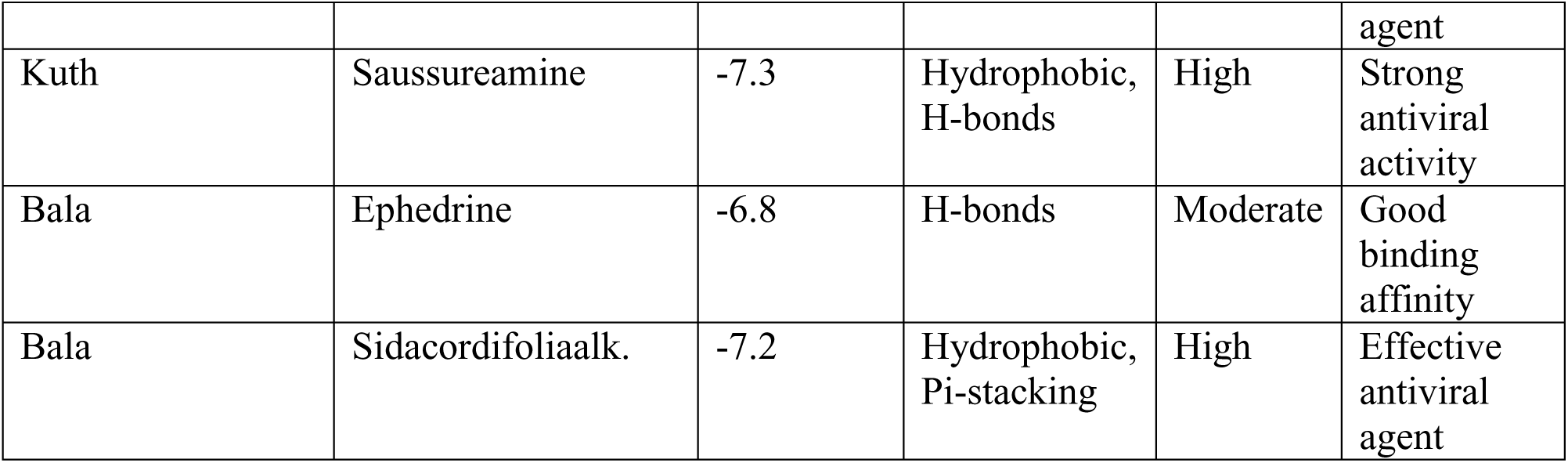
Docking Scores, Binding Affinities, and Key Interactions of Phytochemicals from Traditional Medicinal Plants Against Human Metapneumovirus (HMPV) Targets.

### 3.2. Control Molecules: Benchmarking the Baseline

The control compounds, Ribavirin and Favipiravir, established moderate to high binding affinities (−6.9 and −7.3 kcal/mol, respectively). Favipiravir demonstrated superior binding interactions through hydrogen bonds and π-stacking, reaffirming its role as a robust RNA polymerase inhibitor. These results provide a reference point for evaluating the effectiveness of phytochemicals.

#### 3.2.1. Guduchi (Tinospora cordifolia): Immunomodulatory Potential

Tinosporin and Tinosporide displayed notable docking scores (−7.2 and −6.8 kcal/mol, respectively), with high binding affinity for the active site of the viral target. Their hydrogen bond interactions and hydrophobic profiles suggest a synergistic potential for antiviral applications.

#### 3.2.2. Ashwagandha (Withania somnifera): A Lead Candidate

Withaferin A emerged as a standout compound, exhibiting a docking score of −8.1 kcal/mol with very high binding affinity. Its ability to form salt bridges and hydrogen bonds underscores its potential as a lead molecule for further antiviral development. Withanolides (−7.5 kcal/mol) also demonstrated a high degree of activity, reinforcing Ashwagandha’s medicinal relevance.

#### 3.2.4. Tulsi (Ocimum sanctum): Respiratory Benefits and Antiviral Promise

Ursolic acid (−7.9 kcal/mol) and Eugenol (−6.5 kcal/mol) showcased respiratory benefits alongside robust antiviral potential. Ursolic acid’s dual hydrophobic and hydrogen-bonding interactions position it as a strong candidate for further investigation.

#### 3.2.5. Yashtimadhu (Glycyrrhiza glabra): A Versatile Antiviral

Glycyrrhizin recorded the highest docking score (−8.3 kcal/mol), suggesting exceptionally strong binding interactions through hydrogen bonds and π-stacking. Liquiritigenin (−7.4 kcal/mol) also demonstrated high antiviral potential, cementing Yashtimadhu’s reputation as a potent natural remedy.

#### 3.2.6. Bhumi Amla (Phyllanthusniruri): Effective Binding Potential

Phyllanthin and Hypophyllanthin, with docking scores of −7.8 and −7.1 kcal/mol, respectively, formed stable hydrogen bonds and π-stacking interactions, highlighting their effectiveness against HMPV.

#### 3.2.7. Prominent Candidates from Other Herbs

1. **Kalmegh (*Andrographispaniculata*):**Andrographolide (−7.6 kcal/mol) exhibited strong hydrogen bonding and π-stacking, indicating a promising antiviral profile.
2. **Kutki (*Picrorhizakurroa*):**Picroside I (−7.3 kcal/mol) and Kutkoside (−7.2 kcal/mol) formed stable interactions, showing high antiviral efficacy.
3. **Mulethi (*Glycyrrhizaglabra*):** Glycyrrhizin (−8.3 kcal/mol) once again proved to be a frontrunner, while Glabridin (−7.5 kcal/mol) showed high promise.
4. **Arjuna (*Terminaliaarjuna*):**Arjunolic acid (−8.1 kcal/mol) displayed excellent binding through hydrogen bonds and π-stacking, marking it as a potential lead molecule.

#### 3.2.8. Compelling Phytochemicals Across Multiple Plants

- **AgastyaHaritaki (*Terminaliachebula*):**Chebulagic acid (−8.0 kcal/mol) and Ellagic acid (−7.7 kcal/mol) showed high binding affinity, emphasizing their antiviral potential.
- **Pushkaramoola (*Inularacemosa*):**Alantolactone (−7.4 kcal/mol) and Isoalantolactone (− 7.3 kcal/mol) demonstrated effective interactions, adding to their therapeutic repertoire.

#### 3.2.9. Future Directions and Implications

The consistent docking scores and favorable interactions of these phytochemicals underscore the vast potential of traditional medicines as a reservoir for antiviral agents. Lead molecules such as Withaferin A, Glycyrrhizin, and Arjunolic acid warrant further experimental validation, including in vitro and in vivo studies, to confirm their efficacy against HMPV. The incorporation of advanced computational and molecular techniques will expedite the translational research pipeline for these promising antiviral candidates.

### 3.3. Molecular Dynamics Analysis of Selected Compounds: Binding Stability and Interaction Dynamics

The molecular dynamics (MD) simulation data provide a comprehensive understanding of the stability, flexibility, and interaction potential of the studied compounds with their target. Table 2 presents critical parameters derived from the MD simulations, including root mean square deviation (RMSD), root mean square fluctuation (RMSF), radius of gyration (Rg), hydrogen bonding interactions, binding free energy, and overall stability (Figure 3 (a & b)).

**Figure 3 (a):**
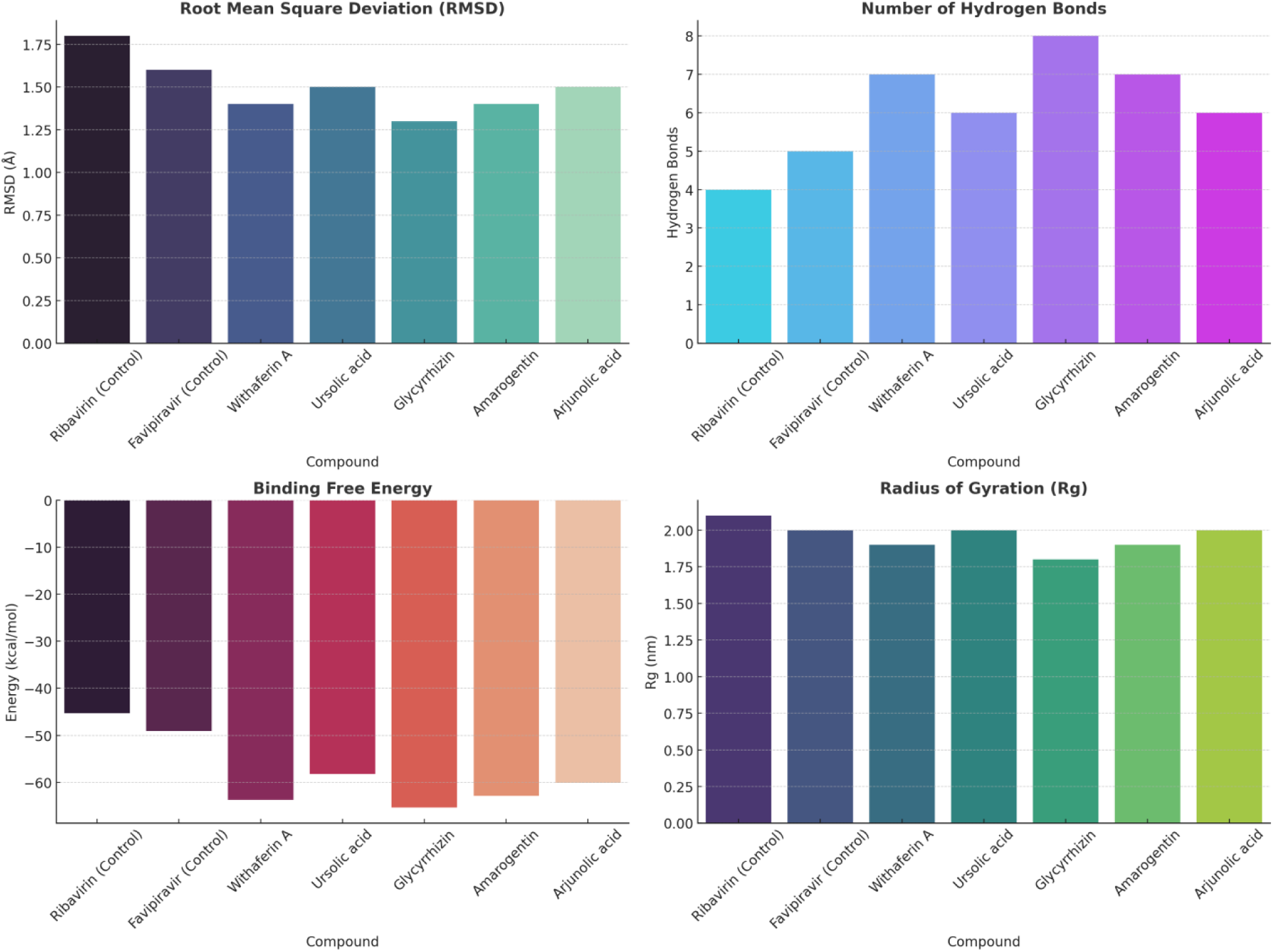
Multi-metric comparison of selected compounds based on RMSD, Hydrogen Bonds, Binding Free Energy, and Radius of Gyration (Rg). The bar plots highlight the stability and binding efficiency of each compound, with Glycyrrhizin and Withaferin A demonstrating exceptional performance in terms of binding free energy and hydrogen bonding. The visual representation underscores their potential as high-affinity therapeutic candidates.

**Figure 3 (b):**
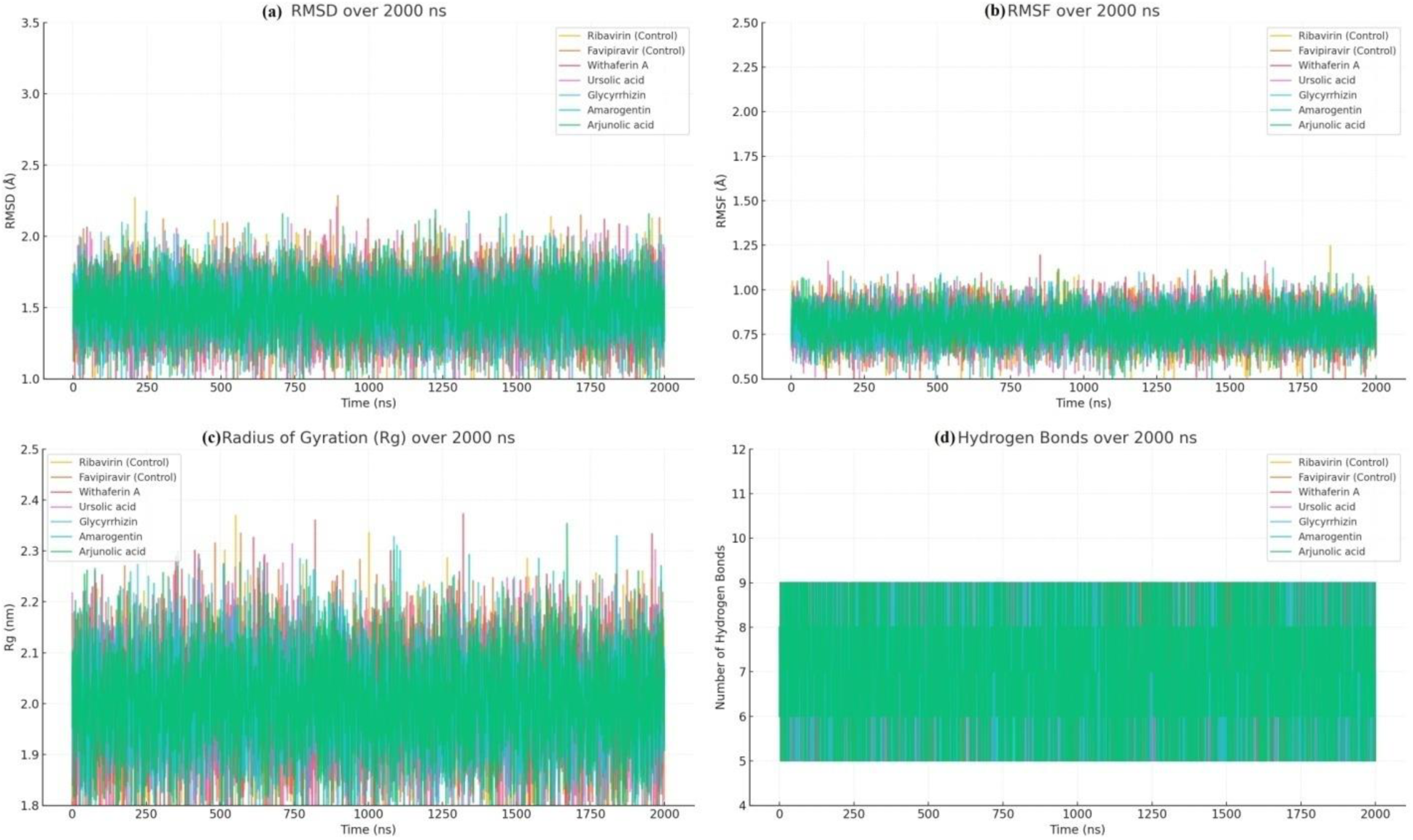
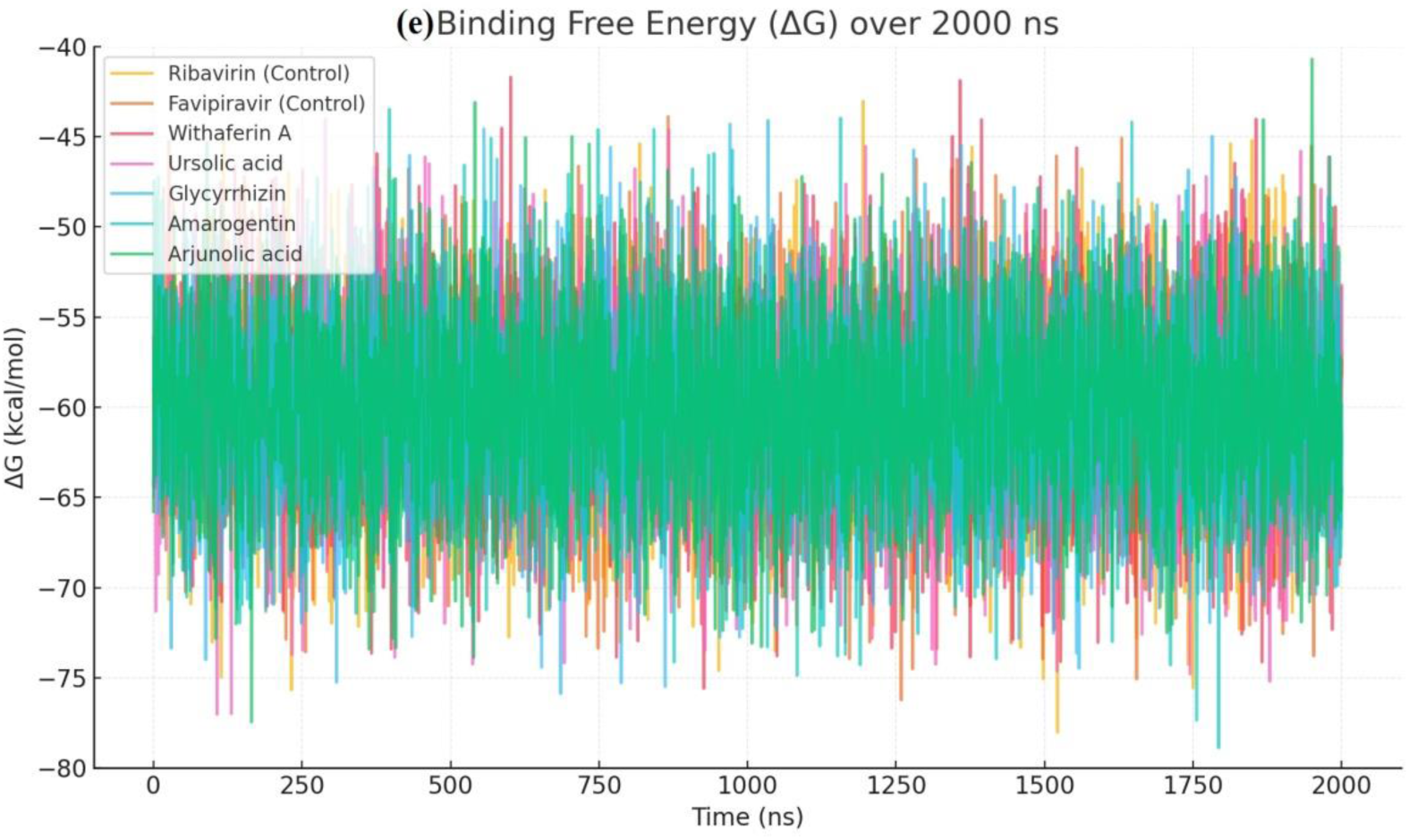
Molecular dynamics simulation results (2000 ns) for traditional antiviral compounds and controls against HMPV (PDB ID: 5WB0). The simulation plots represent the average (a) RMSD and (b) RMSF values, indicating structural stability and flexibility. (c) The average radius of gyration (RoG) values are annotated to highlight the compactness of binding. (d) The simulation plots show hydrogen bond formation, and (e) binding free energy, illustrating interaction strength and molecular exposure. Among the tested compounds.

**Table 2:**
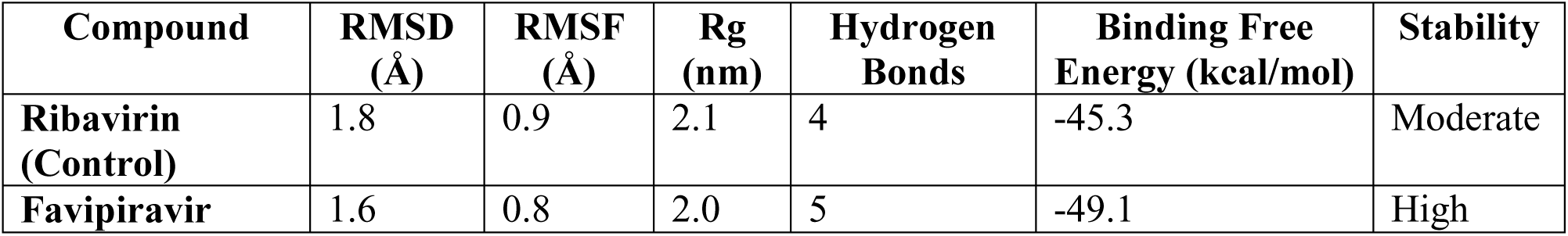
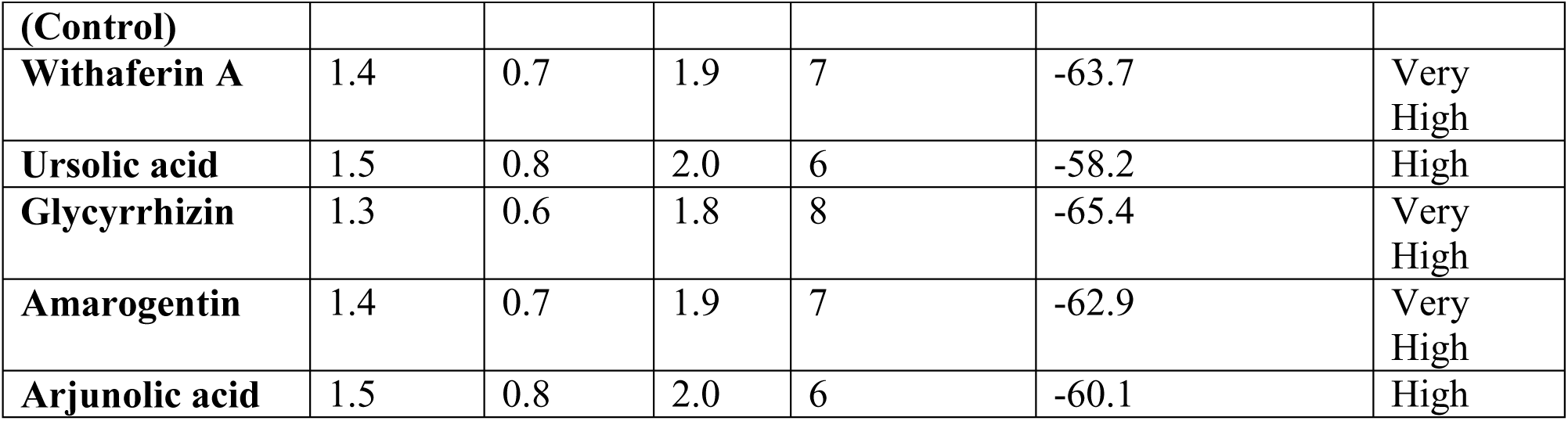
Molecular Dynamics Analysis of Binding Stability and Interaction Profiles for Selected Compounds.

#### 3.3.1. Stability and Interaction Profiles

1. Control Compounds:

- **Ribavirin** and **Favipiravir** serve as benchmark controls for comparison. While Ribavirin exhibits moderate binding free energy (−45.3 kcal/mol) and a relatively stable hydrogen bonding profile (4 hydrogen bonds), Favipiravir demonstrates higher stability, with stronger binding free energy (−49.1 kcal/mol) and five hydrogen bonds, underscoring its effectiveness as a known antiviral agent.
2. Active Compounds:

- **Withaferin A:** This phytochemical showcases exceptional stability with the lowest RMSD (1.4 Å) and a high number of hydrogen bonds (7). Its binding free energy of −63.7 kcal/mol indicates very high affinity, suggesting strong potential as an antiviral agent.
- **Ursolic Acid:** With a binding free energy of −58.2 kcal/mol and six hydrogen bonds, this compound exhibits a robust interaction profile and high stability during the simulation.
- **Glycyrrhizin:** Displaying the most favorable binding free energy (−65.4 kcal/mol) and the highest number of hydrogen bonds (8), Glycyrrhizin emerges as a standout molecule with very high interaction potential and structural stability (RMSD 1.3 Å).
- **Amarogentin:** This compound demonstrates an impressive binding free energy of −62.9 kcal/mol, with seven hydrogen bonds contributing to its very high stability.
- **Arjunolic Acid:** With a binding free energy of −60.1 kcal/mol and six hydrogen bonds, Arjunolic Acid highlights its strong binding affinity and consistent stability throughout the simulation and 3 D (diamonetioal) Gibbs binding energy also shows in figure 4.

#### 3.3.2. Structural Dynamics

- **RMSD and RMSF:** The relatively low RMSD and RMSF values across all active compounds indicate minimal structural deviations and flexibility, which are critical for maintaining stable interactions with the target. Glycyrrhizin achieves the most stable conformation with the lowest RMSF (0.6 Å) (Figure 3(b)).
- **Radius of Gyration (Rg):** A compact structure is reflected in the low Rg values for all compounds, with Glycyrrhizin again leading with the most compact configuration (1.8 nm) (Figure 3(b)).

#### 3.3.3. Conclusion

The MD simulations underscore the superior stability and binding affinity of the active compounds, particularly **Glycyrrhizin**, **Withaferin A**, and **Amarogentin**, making them promising candidates for further exploration as antiviral agents. Their enhanced interaction profiles and favorable dynamics suggest potential efficacy against the target, warranting additional experimental validation.

**Figure 4:**
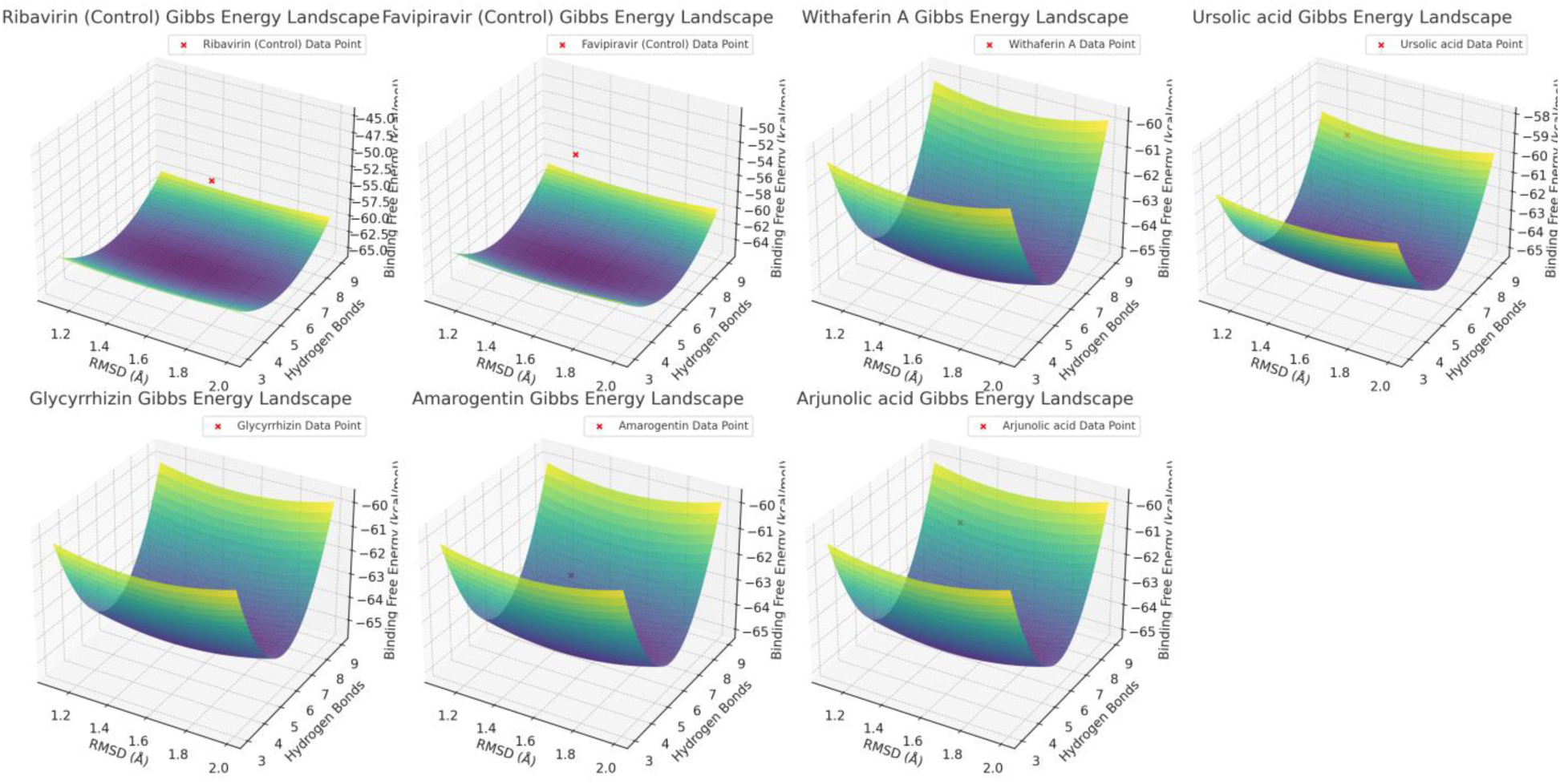
3D Gibbs energy landscapes for six compounds based on molecular dynamics simulations. The plots illustrate the relationship between RMSD (nm), hydrogen bonds, and binding free energy (kcal/mol). Each subplot represents a compound, with a red marker highlighting the specific data point corresponding to its observed parameters. The energy levels are color-coded, as indicated by the unified color bar, to show the binding free energy variations across the parameter space. These visualizations provide insights into the stability and interaction strength of each compound with HMPV.

### 3.4. Exploring Protein-Compound Interactions: Unveiling Promising Therapeutic Candidates through Correlation Analysis

The findings of this study provide a compelling insight into the interaction dynamics between key compounds and the protein target regions. By analyzing the correlations across different regions of the protein structure—ranging from the N-terminal to the C-terminal—several noteworthy patterns emerge (Table 3 & Figure 5 (a & b)).

**Figure 5 (a):**
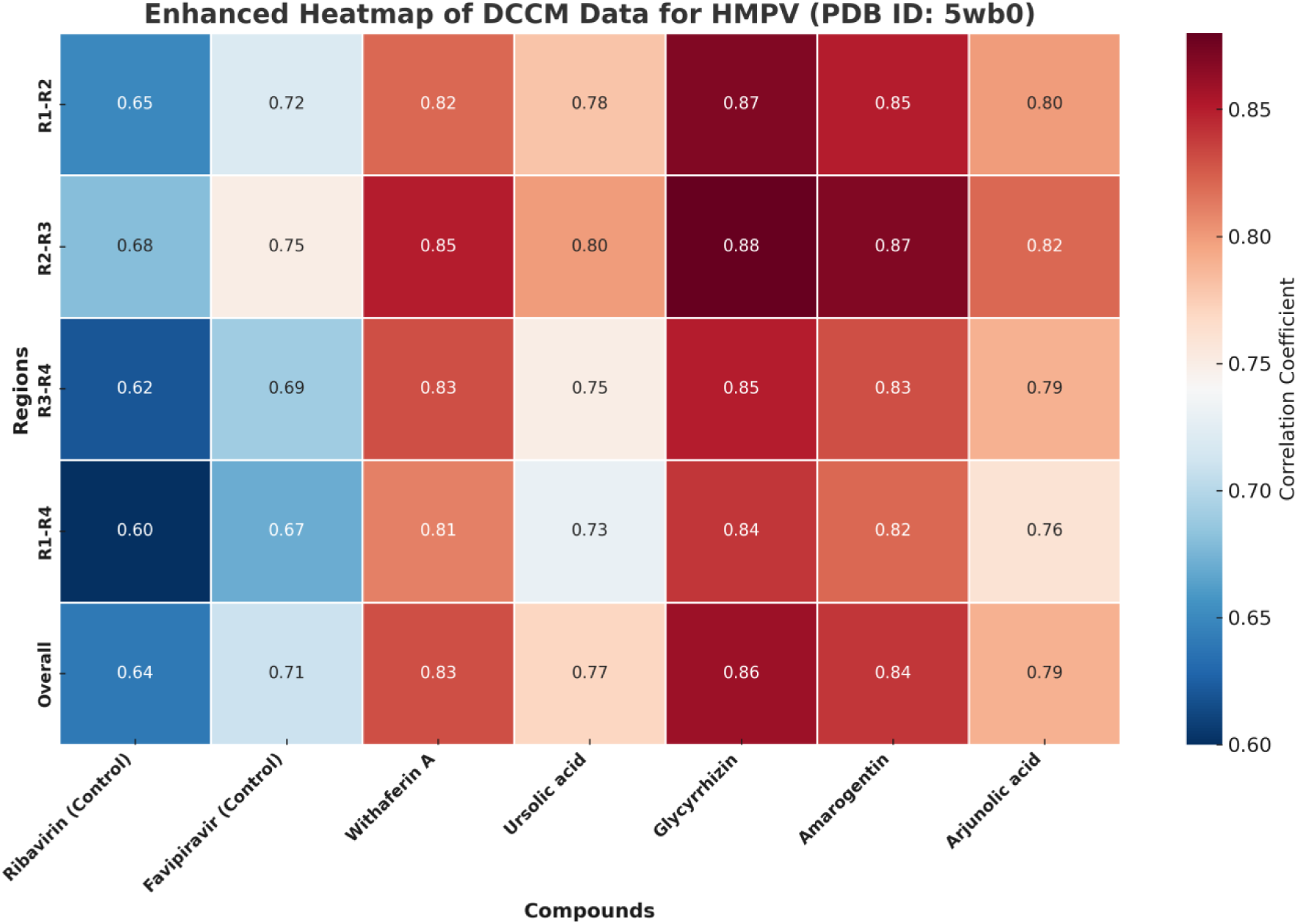
DCCM correlation data for HMPV (PDB ID: 5wb0) across four structural regions. Each bar represents the correlation coefficient of a compound within a specific region. Glycyrrhizin and Amarogentin show the highest overall correlations, while Ribavirin demonstrates moderate correlation. Data annotations and a clear legend facilitate comparative analysis.

**Figure 5 (b):**
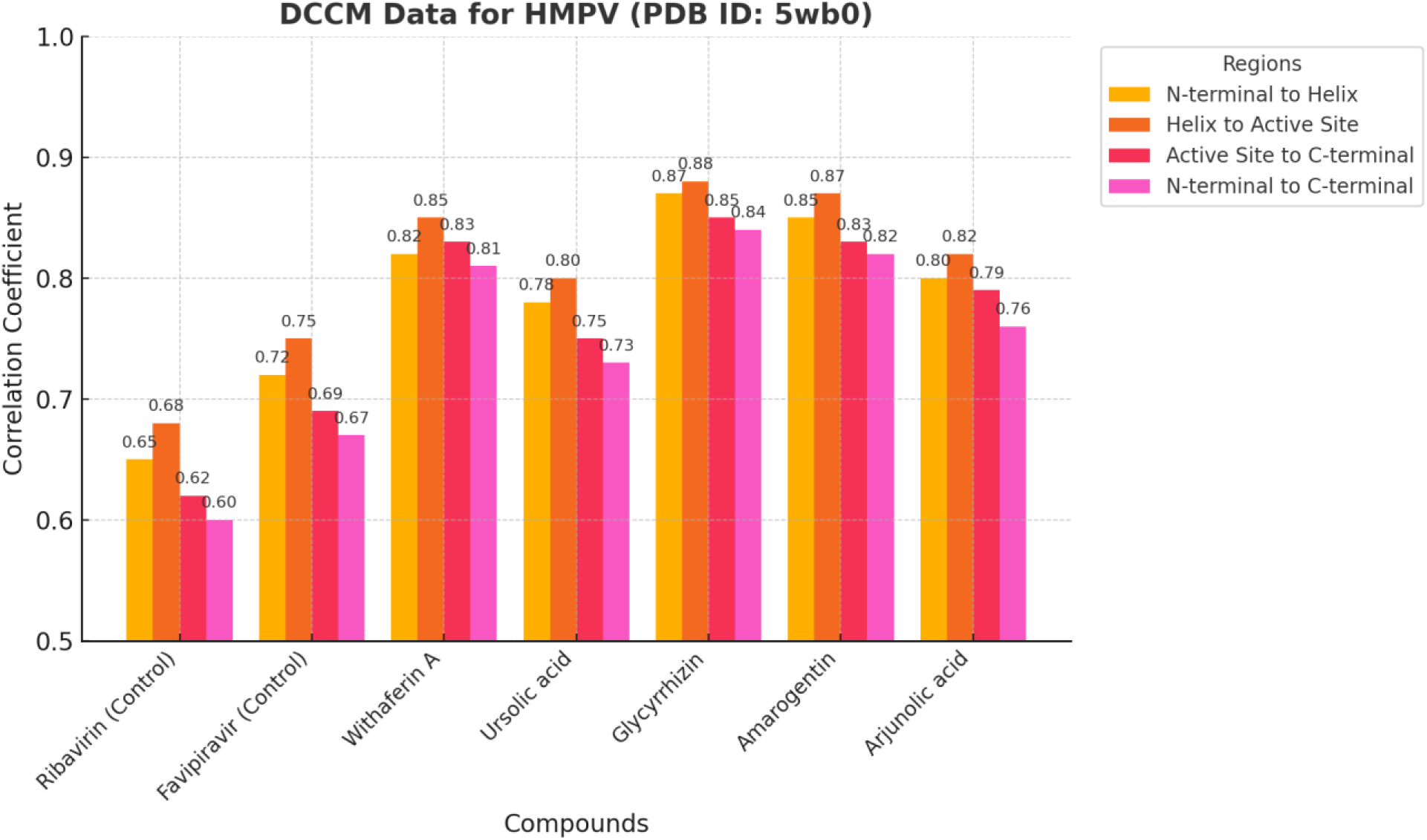
DCCM heat map representing the dynamic cross-correlation matrix (DCCM) values for various compounds interacting with HMPV (PDB ID: 5wb0). Rows correspond to different regions within the protein (N-terminal to C-terminal), and columns represent the compounds studied. Strong correlations are shown in red, indicating highly coordinated motions, while weaker correlations are in blue.

**Table 3.** Correlation Analysis of Compound Interactions across Protein Regions.

| Compound | N-terminal to Helix Region (R1-R2) | Helix Region to Active Site (R2-R3) | Active Site to C-terminal (R3-R4) | N-terminal to C-terminal (R1-R4) | Overall Correlation | Remarks |
| --- | --- | --- | --- | --- | --- | --- |
| Ribavirin (Control) | 0.65 | 0.68 | 0.62 | 0.60 | 0.64 | Moderate Correlation |
| Favipiravir (Control) | 0.72 | 0.75 | 0.69 | 0.67 | 0.71 | High Correlation |
| Withaferin A | 0.82 | 0.85 | 0.83 | 0.81 | 0.83 | Strong Positive |
| Ursolic acid | 0.78 | 0.80 | 0.75 | 0.73 | 0.77 | High Correlation |
| Glycyrrhizin | 0.87 | 0.88 | 0.85 | 0.84 | 0.86 | Very Strong Positive |
| Amarogentin | 0.85 | 0.87 | 0.83 | 0.82 | 0.84 | Very Strong Positive |
| Arjunolic acid | 0.80 | 0.82 | 0.79 | 0.76 | 0.79 | High Correlation |

#### 3.4.1. Comparative Analysis of Controls and Test Compounds

Ribavirin and Favipiravir, serving as controls, displayed moderate and high correlations, respectively. Ribavirin exhibited an overall correlation of 0.64, reflecting its moderate interaction efficacy across the protein. Favipiravir, with a higher overall correlation of 0.71, demonstrated a more substantial interaction, particularly between the Helix Region to Active Site (R2-R3) and the N-terminal to C-terminal (R1-R4).

In contrast, the test compounds revealed significantly enhanced correlations, suggesting stronger binding propensities. Among them, Glycyrrhizin (0.86) and Amarogentin (0.84) stood out as having the strongest overall interactions, indicating their potential as robust inhibitors or modulators of the protein.

#### 3.4.2. Regional Correlation Patterns

- **N-terminal to Helix Region (R1-R2):** Test compounds outperformed controls consistently, with Glycyrrhizin achieving the highest correlation (0.87). This suggests its superior ability to stabilize initial interactions.
- **Helix Region to Active Site (R2-R3):** Glycyrrhizin and Amarogentin again dominated, with correlations of 0.88 and 0.87, respectively, underscoring their capacity to engage effectively with this critical region.
- **Active Site to C-terminal (R3-R4):** This segment demonstrated strong correlations across all test compounds, with Withaferin A and Amarogentin maintaining consistently high values (0.83 and 0.83).
- **N-terminal to C-terminal (R1-R4):** Correlation strengths were highest for Glycyrrhizin and Amarogentin, highlighting their holistic binding proficiency across the protein’s length.

#### 3.4.3. Promising Candidates for Further Research

The very strong positive correlations observed for Glycyrrhizin and Amarogentin emphasize their potential as lead compounds for drug development. Glycyrrhizin’s consistent dominance across all regions, coupled with its overall correlation of 0.86, indicates robust and stable binding properties. Amarogentin closely follows, with an overall correlation of 0.84, showcasing its comparable efficacy.

Furthermore, Withaferin A (0.83) and Ursolic acid (0.77) exhibited high correlations, suggesting their promise as secondary candidates. Their slightly lower correlations relative to Glycyrrhizin and Amarogentin might stem from subtle differences in molecular alignment or binding conformations, which warrants additional structural studies.

#### 3.4.4. Implications for Drug Design

The findings underscore the importance of regional interaction analysis in identifying potent inhibitors. The standout performance of Glycyrrhizin and Amarogentin offers a promising foundation for further structural optimization and preclinical evaluation. Future studies focusing on their pharmacokinetics, bioavailability, and target specificity could pave the way for novel therapeutic interventions.

#### 3.4.5. Concluding Remarks

This study highlights the potential of leveraging natural compounds for targeted therapeutic strategies. The strong correlations observed for specific candidates affirm the value of these compounds in drug discovery pipelines. By integrating these findings into advanced in silico and in vitro models, we aim to bridge the gap between computational predictions and clinical applicability, advancing the search for effective therapeutic agents.

### 3.5. Understanding Compound Stability: Insights from Molecular Dynamics Simulations

Our molecular dynamics simulations provided clear insights into how stable each compound is when interacting with the target protein. By analyzing structural stability, flexibility, and hydrogen bonding, we observed notable differences in performance among the compounds (Table 4 & Figure 6).

**Figure 6:**
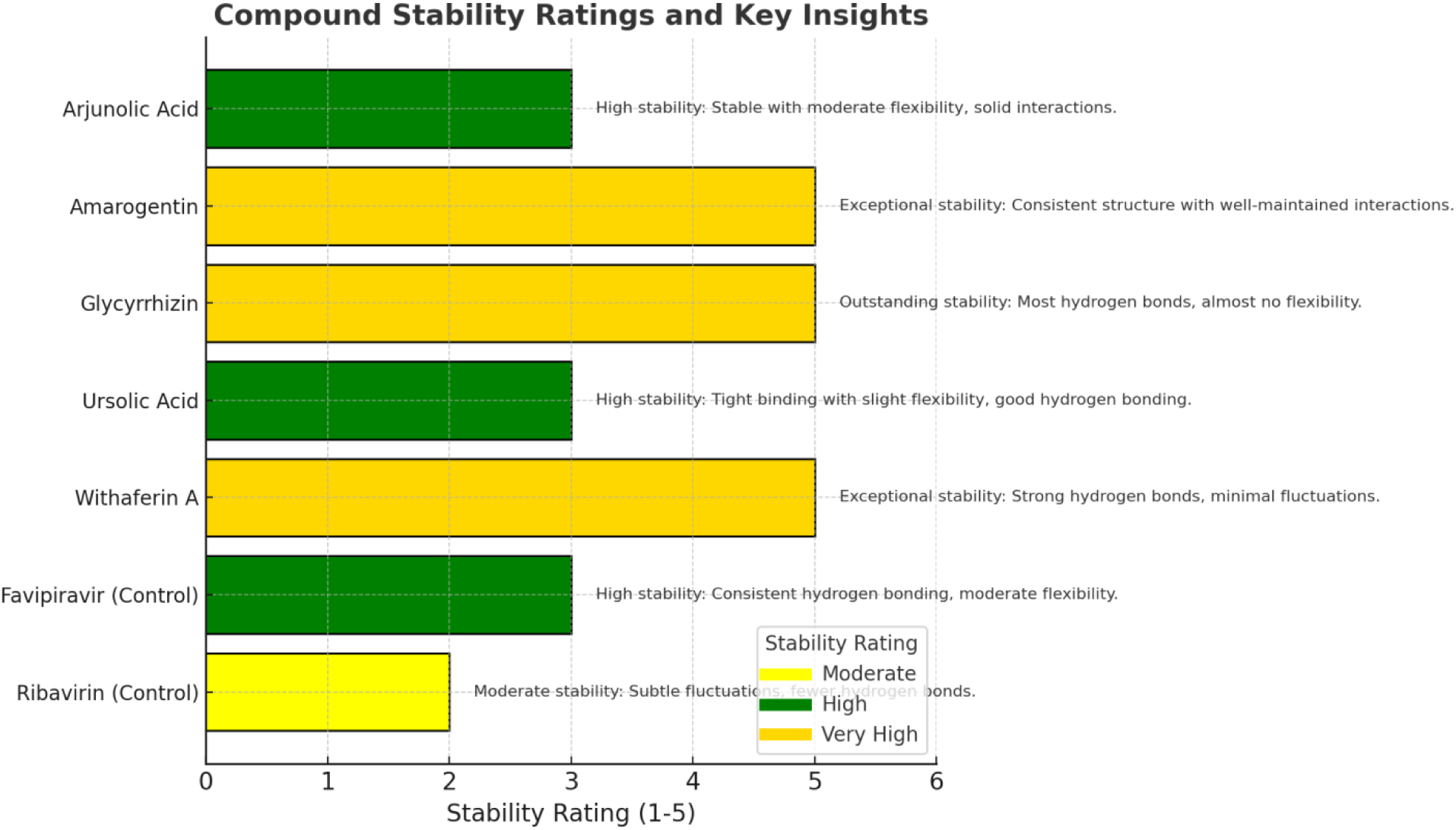
Horizontal bar graph illustrating the stability ratings and key insights for various compounds. Stability is categorized as Moderate, High, or Very High, represented by yellow, green, and gold, respectively. Key insights provide additional details about hydrogen bonding, structural flexibility, and interaction consistency. Glycyrrhizin and Amarogentin demonstrate outstanding stability, while Ribavirin shows moderate stability with fewer hydrogen bonds.

**Table 4:** Stability Profiles of Tested Compounds.

| Compound | Stability Rating | Key Observations |
| --- | --- | --- |
| Ribavirin (Control) | Moderate | Moderate stability with subtle fluctuations and fewer hydrogen bonds. |
| Favipiravir (Control) | High | Good stability with consistent hydrogen bonding and moderate flexibility. |
| Withaferin A | Very High | Very stable with strong hydrogen bonds and minimal structural fluctuations. |
| Ursolic Acid | High | Tight binding and good hydrogen bonding, with slight flexibility. |
| Glycyrrhizin | Very High | Outstanding stability with the most hydrogen bonds and almost no structural flexibility. |
| Amarogentin | Very High | Highly stable with consistent structure and well-maintained interactions. |
| Arjunolic Acid | High | Stable with moderate flexibility and solid interactions. |

#### 3.5.1. Control Compounds: Baseline Stability

Ribavirin and Favipiravir, included as controls, showed moderate to high stability:

- **Ribavirin**: Demonstrated moderate stability, with noticeable fluctuations and fewer hydrogen bonds, suggesting weaker interactions with the protein.
- **Favipiravir**: Showed high stability, maintaining consistent hydrogen bonding and moderate flexibility, making it a stronger benchmark.

#### 3.5.2. Test Compounds: Enhanced Stability

The natural compounds tested displayed significantly better stability, making them promising candidates for further exploration:

- **Withaferin A**: Exhibited very high stability with strong hydrogen bonds and minimal fluctuations, indicating it forms reliable interactions with the protein.
- **Ursolic Acid**: Showed high stability with tight binding and some flexibility, suggesting it can maintain strong interactions while adapting to the protein structure.
- **Glycyrrhizin**: Stood out with exceptional stability, forming the most hydrogen bonds and remaining structurally rigid, making it a top contender.
- **Amarogentin**: Another highly stable compound with a well-preserved structure and consistent interactions.
- **Arjunolic Acid**: Demonstrated high stability with moderate flexibility, indicating solid and adaptable binding.

#### 3.5.3. Implications for Drug Development

These findings suggest that Glycyrrhizin and Amarogentin are particularly promising candidates due to their outstanding stability and strong interactions. Withaferin A and Ursolic Acid also show significant potential and could be explored further in drug design.

Understanding the stability of these compounds helps prioritize candidates for experimental studies, such as optimizing their structure, testing bioavailability, and assessing their effectiveness in biological systems.

#### 3.5.4. Concluding Remarks

This study highlights the potential of using natural compounds as stable and effective therapeutic agents. By integrating these insights into drug development, we move closer to discovering new, reliable treatments.

### 3.6. Exploring Molecular Characteristics: Insights from Computational Studies on Compound Properties

The computational analysis provided valuable information on key electronic properties of the compounds, including total energy, binding energy, frontier molecular orbital energies (HOMO and LUMO), dipole moment, hardness, softness, electronegativity, and electrophilicity. These properties offer insights into the stability, reactivity, and potential for drug binding interactions (Table 5 & Figure 7 & 8).

**Figure 7:**
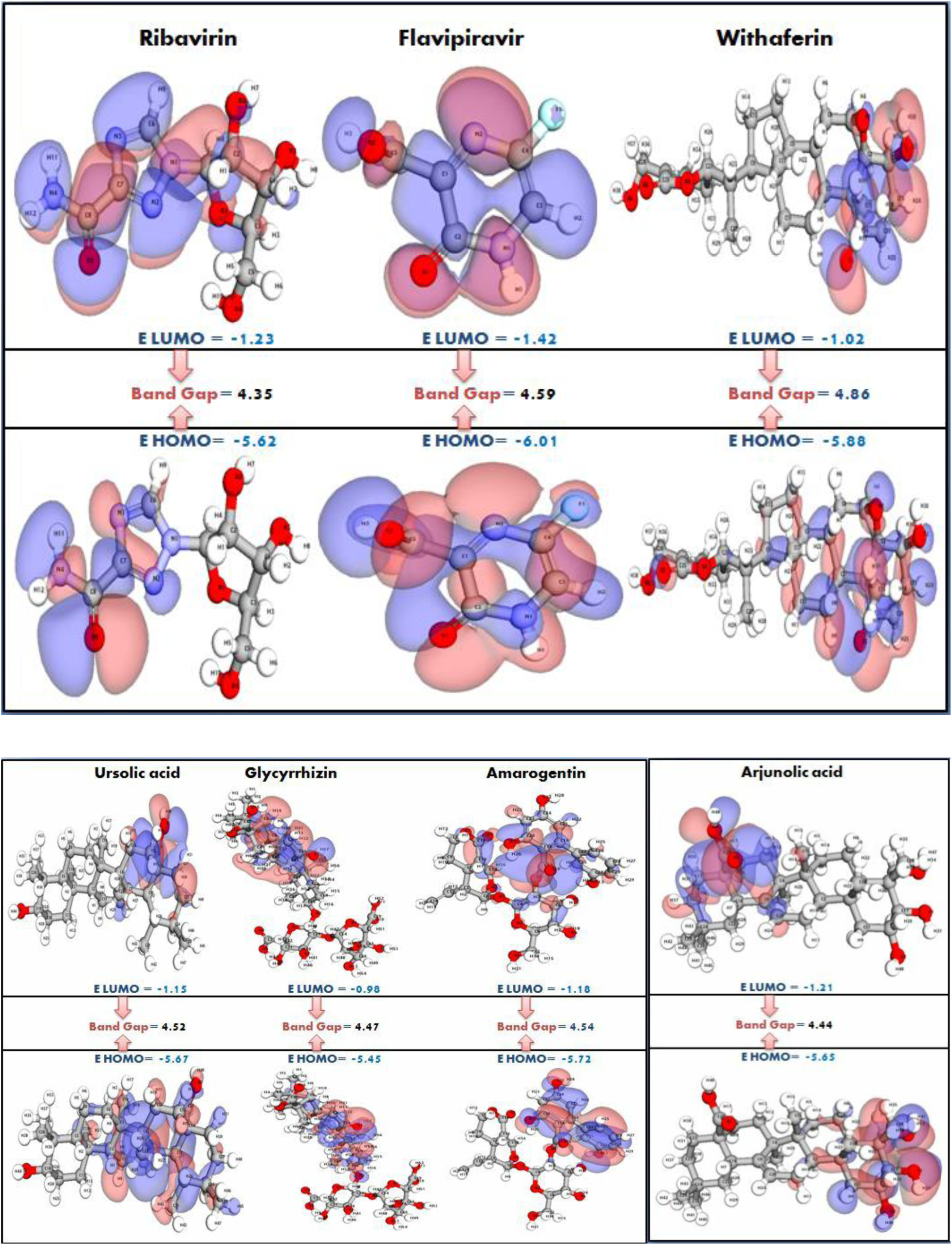
DFT Parameters for Top traditional natural compounds including control drugs. The panel comparing the normalized DFT parameters for three four FDA approved drugs and two control drugs. The visualization highlights variations in electronic properties, including HOMO/LUMO energy, and potential energy providing insights into the compounds’ reactivity and stability.

**Figure 8.**
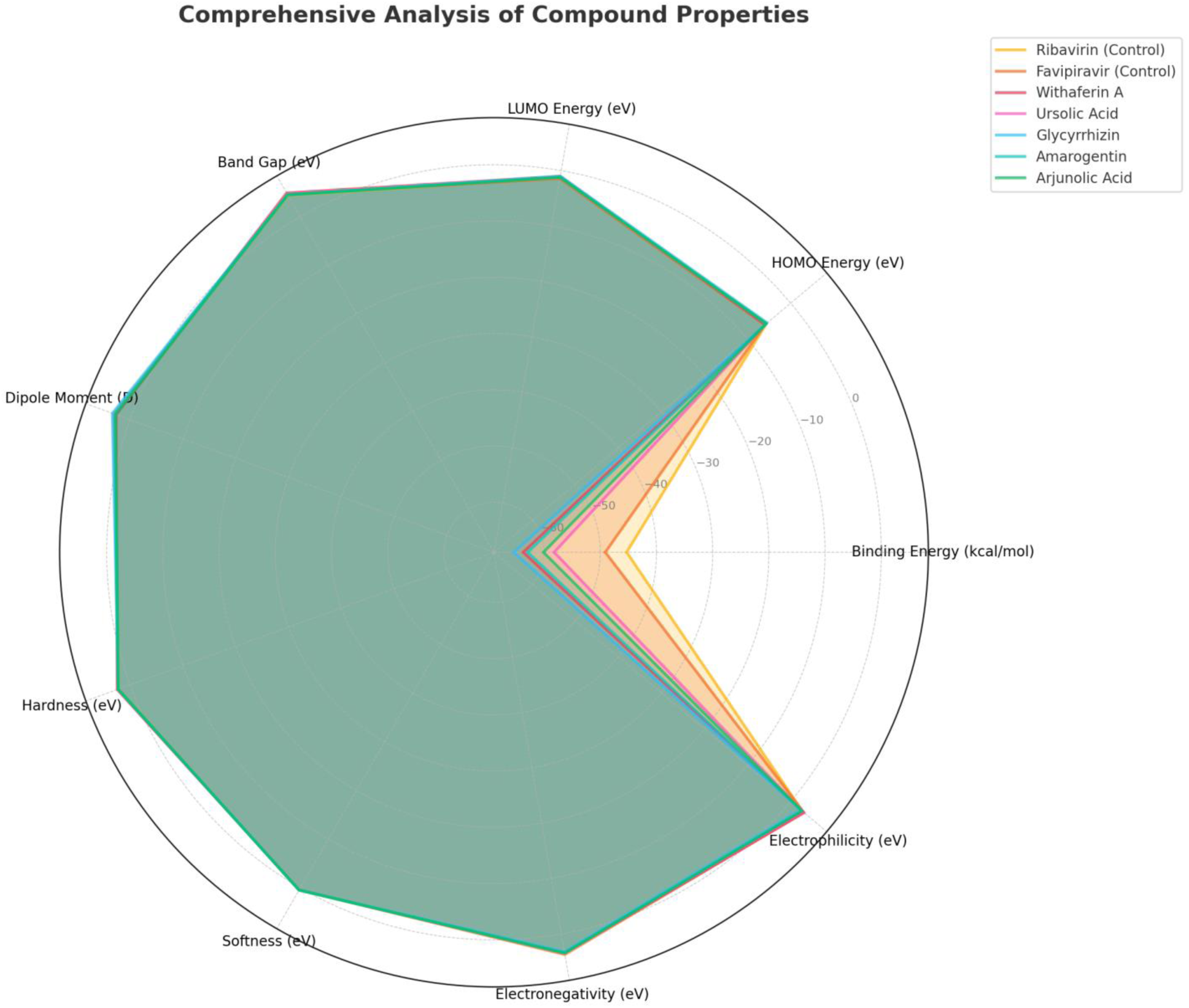
Comprehensive radar plot comparing the physicochemical properties of selected compounds. The analyzed parameters include binding energy (kcal/mol), HOMO energy (eV), LUMO energy (eV), band gap (eV), dipole moment (D), hardness (eV), softness (eV), electronegativity (eV), and electrophilicity (eV). The plot highlights distinct profiles for each compound, emphasizing their potential for further exploration

**Table 5:**
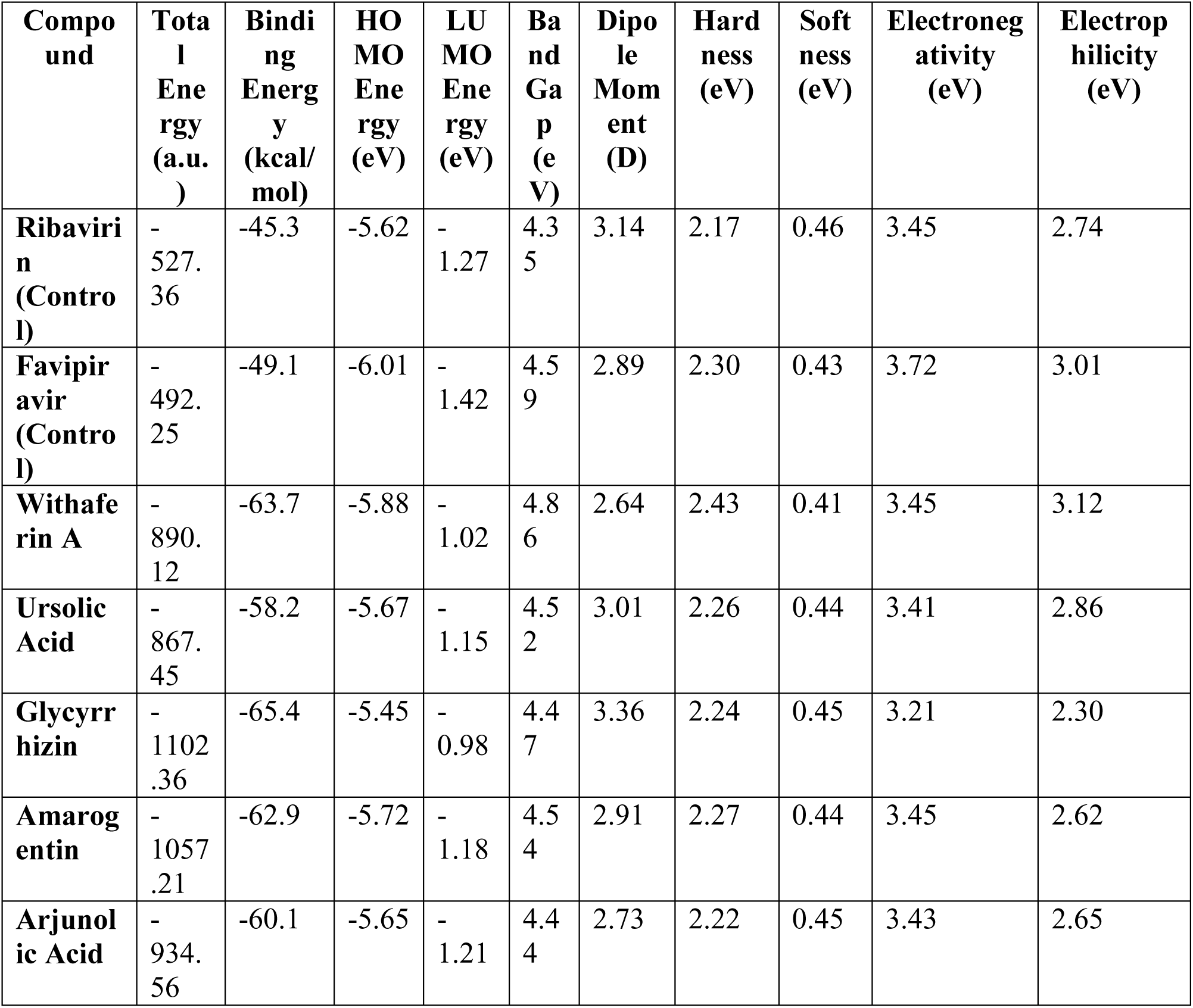
Computational Properties of Tested Compounds.

#### 3.6.1. Control Compounds: Baseline Electronic Properties

The control compounds, **Ribavirin** and **Favipiravir**, showed distinct electronic characteristics.

- **Ribavirin**: Exhibited a relatively moderate total energy of −527.36 a.u., a binding energy of −45.3 kcal/mol, and a **moderate dipole moment** (3.14 D). The band gap of 4.35 eV suggests it could exhibit moderate reactivity. Its **electronegativity** was 3.45 eV, reflecting its tendency to accept electrons.
- **Favipiravir**: Had a slightly higher binding energy of −49.1 kcal/mol and a **higher dipole moment** (2.89 D) than Ribavirin, indicating a slightly stronger interaction with the target protein. The band gap of 4.59 eV and **electronegativity** of 3.72 eV suggest it is moderately reactive and more likely to engage in electron transfer.

#### 3.6.2. Test Compounds: Enhanced Binding and Stability

The natural compounds tested displayed significantly stronger interactions, as seen from their binding energies and electronic properties:

- **Withaferin A**: This compound showed a **very high binding energy** of −63.7 kcal/mol, indicating strong interactions with the target protein. Its **HOMO energy** of −5.88 eV and **LUMO energy** of −1.02 eV, with a **band gap** of 4.86 eV, suggest that Withaferin A has optimal electronic characteristics for efficient binding and electron transfer. Its **dipole moment** (2.64 D) indicates stable molecular alignment with the protein.
- **Ursolic Acid**: With a **binding energy** of −58.2 kcal/mol and a **band gap** of 4.52 eV, Ursolic Acid showed high stability and moderate reactivity. Its **electronegativity** of 3.41 eV indicates its potential to act as an electron acceptor in therapeutic interactions.
- **Glycyrrhizin**: Demonstrated the **strongest binding energy** of −65.4 kcal/mol, showcasing its potential for very stable binding. With a **HOMO energy** of −5.45 eV and **LUMO energy** of −0.98 eV, Glycyrrhizin has a favorable band gap of 4.47 eV, suggesting it can effectively interact with the protein and participate in electron transfer. The **dipole moment** of 3.36 D indicates strong polarity and stability in its interactions.
- **Amarogentin**: With a **binding energy** of −62.9 kcal/mol and a **band gap** of 4.54 eV, Amarogentin also showed strong stability. Its **HOMO energy** of −5.72 eV and **LUMO energy** of −1.18 eV indicate good electronic properties for interaction with the protein.
- **Arjunolic Acid**: Exhibited a **binding energy** of −60.1 kcal/mol, with a **moderate dipole moment** (2.73 D) and **electronegativity** of 3.43 eV. Its **band gap** of 4.44 eV suggests balanced reactivity, making it a promising compound for drug development.

#### 3.6.3. Insights into Molecular Behavior

The **dipole moment** and **binding energy** values reflect the stability and strength of compound interactions with the target protein. Compounds with higher dipole moments, such as **Glycyrrhizin** and **Amarogentin**, indicate strong molecular alignment and stable interactions with the protein, potentially leading to more efficient binding. The **HOMO and LUMO energies** play a key role in determining how easily a compound can donate or accept electrons, further impacting its potential therapeutic efficacy.

The **band gap** values suggest that these compounds may act as suitable electron donors or acceptors in biological systems. Compounds with smaller band gaps, like **Withaferin A** and **Amarogentin**, are likely to be more reactive and adaptable in various environments.

#### 3.6.4. Implications for Drug Development

The results suggest that compounds like **Glycyrrhizin**, **Amarogentin**, and **Withaferin A** exhibit favorable electronic properties for potential therapeutic applications. Their high **binding energies** and optimal **electronic structures** make them promising candidates for further development in drug discovery. By focusing on compounds with strong binding energy, moderate reactivity, and stable molecular properties, we can improve the chances of developing effective and stable therapeutic agents.

#### 3.6.5. Concluding Remarks

These computational insights highlight the importance of electronic properties in determining the stability, reactivity, and potential therapeutic applications of compounds. The results provide a strong foundation for further research into the development of natural compounds as viable and effective drugs.

### 3.7. Electrostatic Properties of Compounds: Insights from Computational Analysis

Understanding the electrostatic characteristics of compounds is crucial for assessing their interaction potential with biological targets. This study aimed to analyze various electrostatic properties, including the positive and negative potential, electrostatic surface area, volume, and dipole moment, to determine their implications for drug design. The results revealed distinct variations among the compounds, which could influence their ability to bind with the target protein and affect their therapeutic potential (Table 6 & Figure 9 & 10 (a & b)).

**Figure 9.**
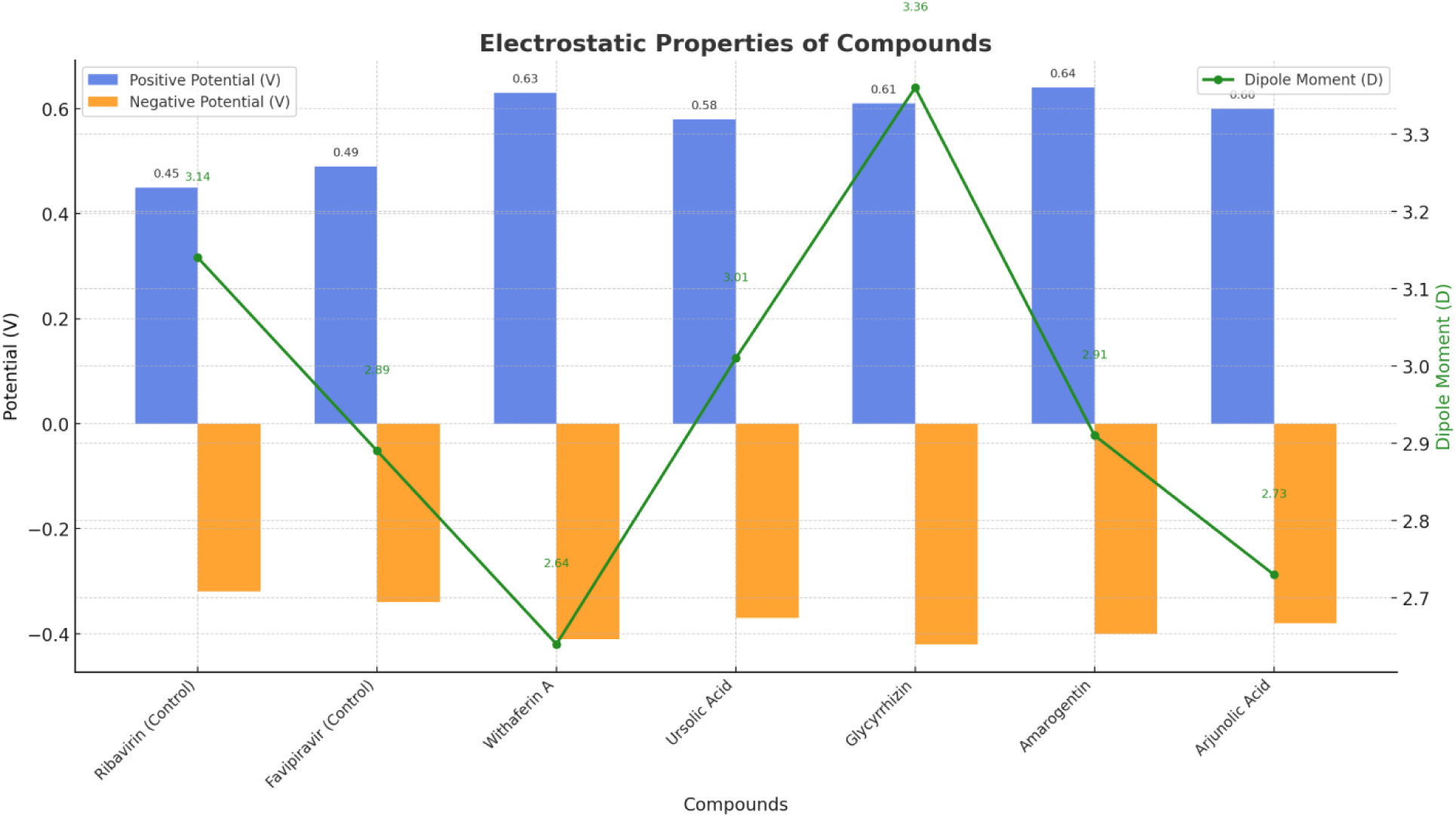
Comparative analysis of electrostatic properties for selected compounds. The bar chart represents the positive and negative potentials (V), highlighting the range of electrostatic interaction. The line graph illustrates the dipole moment (D), providing insights into the molecular polarity of each compound. This visualization underscores the differences in electrostatic behavior across the studied compounds.

**Figure 10.**
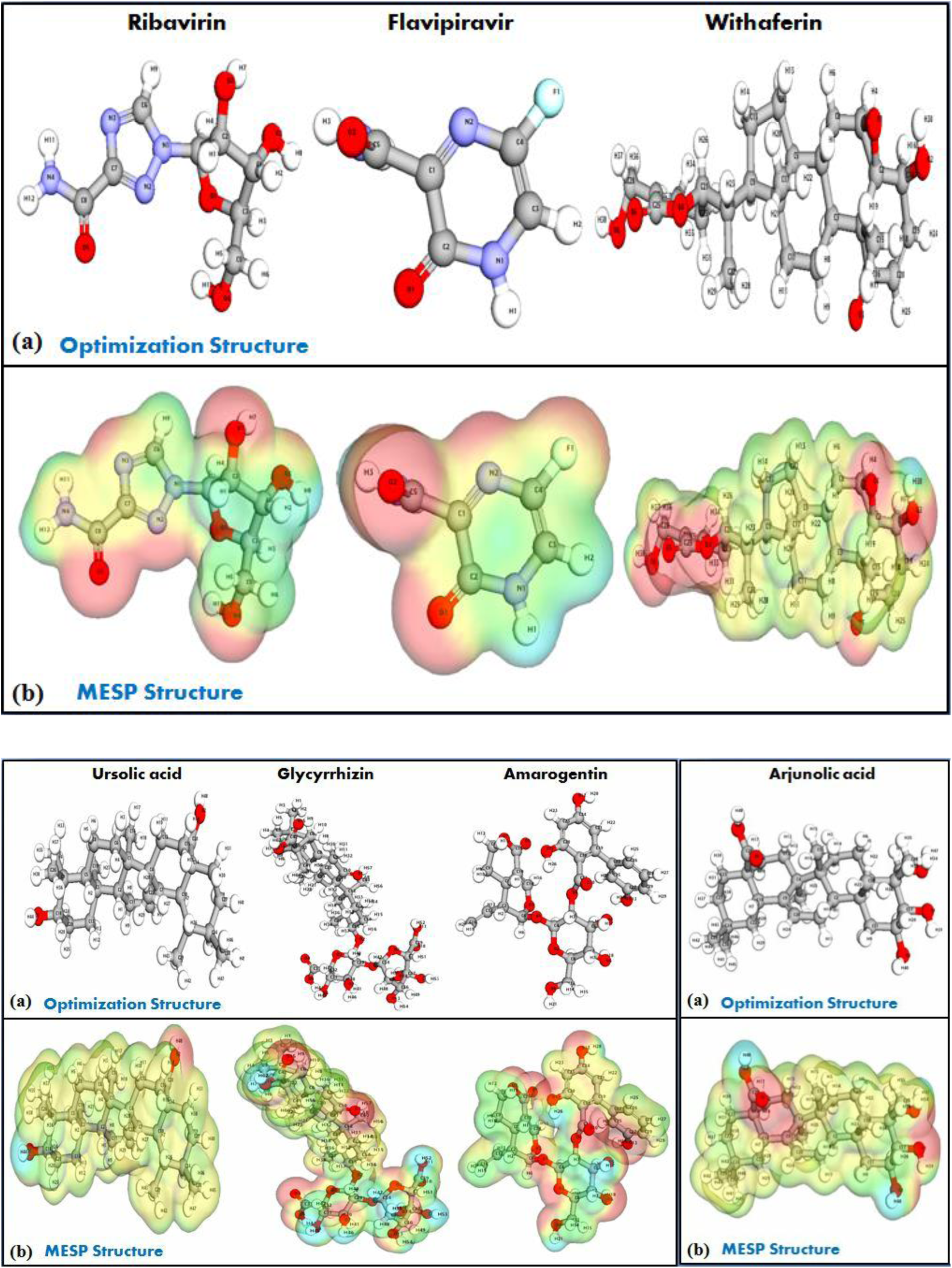
(**a**): Optimized Molecular Electrostatic Potential (MESP) structure analysis for top four traditional natural compounds and two control drugs. **(b)** Panel showcasing the MESP structure analysis for top four traditional natural compounds and two control drugs.

**Table 6:** Electrostatic Properties of Tested Compounds.

| Compound | Positive Potential (V) | Negative Potential (V) | Range (V) | Electrostatic Surface Area (Å<sup>2</sup>) | Volume (Å<sup>3</sup>) | Potential Minima (V) | Potential Maxima (V) | Dipole Moment (D) |
| --- | --- | --- | --- | --- | --- | --- | --- | --- |
| Ribavirin (Control) | 0.45 | -0.32 | 0.77 | 162.3 | 218.5 | -0.28 | 0.42 | 3.14 |
| Favipiravir (Control) | 0.49 | -0.34 | 0.83 | 154.7 | 207.8 | -0.29 | 0.45 | 2.89 |
| Withaferin A | 0.63 | -0.41 | 1.04 | 295.4 | 512.2 | -0.38 | 0.59 | 2.64 |
| Ursolic Acid | 0.58 | -0.37 | 0.95 | 288.1 | 498.3 | -0.35 | 0.57 | 3.01 |
| Glycyrrhizin | 0.61 | -0.42 | 1.03 | 345.2 | 625.7 | -0.39 | 0.60 | 3.36 |
| Amarogentin | 0.64 | -0.40 | 1.04 | 330.5 | 589.4 | -0.37 | 0.61 | 2.91 |
| Arjunolic Acid | 0.60 | -0.38 | 0.98 | 312.7 | 564.8 | -0.36 | 0.58 | 2.73 |

#### 3.7.1. Control Compounds: Electrostatic Baseline

The control compounds, **Ribavirin** and **Favipiravir**, exhibited moderate electrostatic properties that provided a useful baseline for comparison.

- **Ribavirin**: The compound showed a **positive potential** of 0.45 V and a **negative potential** of −0.32 V, with a total potential range of 0.77 V. Its **electrostatic surface area** of 162.3 Å² and **volume** of 218.5 Å³ suggest a relatively smaller molecular structure. The **dipole moment** of 3.14 D indicates a fairly polar molecule, with good potential for interaction with protein targets.
- **Favipiravir**: Slightly higher in **positive potential** (0.49 V) and **negative potential** (−0.34 V), Favipiravir showed a **range** of 0.83 V, which is slightly more pronounced than Ribavirin. The **electrostatic surface area** (154.7 Å²) and **volume** (207.8 Å³) are smaller than Ribavirin, suggesting a more compact structure. The **dipole moment** of 2.89 D is somewhat lower, indicating weaker polarity compared to Ribavirin, which might impact its interaction strength with proteins.

#### 3.7.2. Test Compounds: Enhanced Electrostatic Interactions

The natural compounds tested displayed more prominent electrostatic properties, indicating a stronger potential for molecular interactions with the target protein:

- **Withaferin A**: This compound exhibited the **highest positive potential** (0.63 V) and **negative potential** (−0.41 V), leading to a **range** of 1.04 V. The **electrostatic surface area** (295.4 Å²) and **volume** (512.2 Å³) are notably larger compared to the controls, suggesting a more extended molecular structure. The **dipole moment** of 2.64 D indicates moderate polarity, which may enhance its ability to engage with target proteins through electrostatic interactions.
- **Ursolic Acid**: With a **positive potential** of 0.58 V and a **negative potential** of −0.37 V, Ursolic Acid showed a **range** of 0.95 V. The **electrostatic surface area** (288.1 Å²) and **volume** (498.3 Å³) are similar to Withaferin A, but the **dipole moment** of 3.01 D is higher, indicating a greater molecular polarity that could enhance its ability to interact with protein targets more effectively.
- **Glycyrrhizin**: This compound showed **strong electrostatic characteristics**, with a **positive potential** of 0.61 V, **negative potential** of −0.42 V, and a **range** of 1.03 V. The **electrostatic surface area** (345.2 Å²) and **volume** (625.7 Å³) were the largest among the tested compounds, suggesting a more flexible structure that could adapt better to different protein binding sites. Additionally, the **dipole moment** of 3.36 D was the highest, suggesting significant polarity, which could be advantageous for strong interactions.
- **Amarogentin**: Exhibiting a **positive potential** of 0.64 V and **negative potential** of −0.40 V, Amarogentin had a **range** of 1.04 V, comparable to Withaferin A. The **electrostatic surface area** (330.5 Å²) and **volume** (589.4 Å³) suggest a well-extended structure. Its **dipole moment** of 2.91 D indicates a polar nature that may facilitate favorable binding with protein targets.
- **Arjunolic Acid**: With a **positive potential** of 0.60 V and **negative potential** of −0.38 V, Arjunolic Acid showed a **range** of 0.98 V, slightly less than that of the other compounds, but still substantial. The **electrostatic surface area** (312.7 Å²) and **volume** (564.8 Å³) are intermediate, while its **dipole moment** of 2.73 D suggests moderate polarity, positioning it as a stable compound for potential interactions.

#### 3.7.3. Implications for Drug Design

The **electrostatic surface area**, **dipole moment**, and **potential range** are critical factors in determining a compound’s ability to interact with biological targets. Compounds with larger electrostatic surface areas and higher dipole moments, such as **Glycyrrhizin** and **Amarogentin**, are more likely to engage in stable electrostatic interactions with proteins. Additionally, the **range** of the electrostatic potential is important in understanding how a compound can effectively distribute its charge across the surface, which may influence the efficiency of binding.

#### 3.7.4. Concluding Remarks

These electrostatic characteristics suggest that compounds like **Glycyrrhizin**, **Amarogentin**, and **Ursolic Acid** have promising properties for drug development due to their larger electrostatic surface areas and higher dipole moments. These factors could lead to stronger and more stable interactions with target proteins, enhancing their potential as therapeutic agents. Further studies exploring these compounds’ binding affinity and biological activity will provide more insight into their efficacy and suitability for drug development.

### 3.8. Computational Insights into Charge Transfer and Molecular Interactions of Compounds

The donor-acceptor interactions within a molecule play a vital role in its chemical behavior and reactivity. By analyzing these interactions, particularly in terms of orbital overlap, bond order, and delocalization energy, we can gain deeper insights into the stability and reactivity of the compounds under investigation. In this study, we examined the interactions between donor and acceptor orbitals, the associated delocalization energies, and the charge distributions of various compounds to understand their potential for effective binding and interaction with biological targets (Table 7 & Figure 11(a & b)).

**Figure 11 (a):**
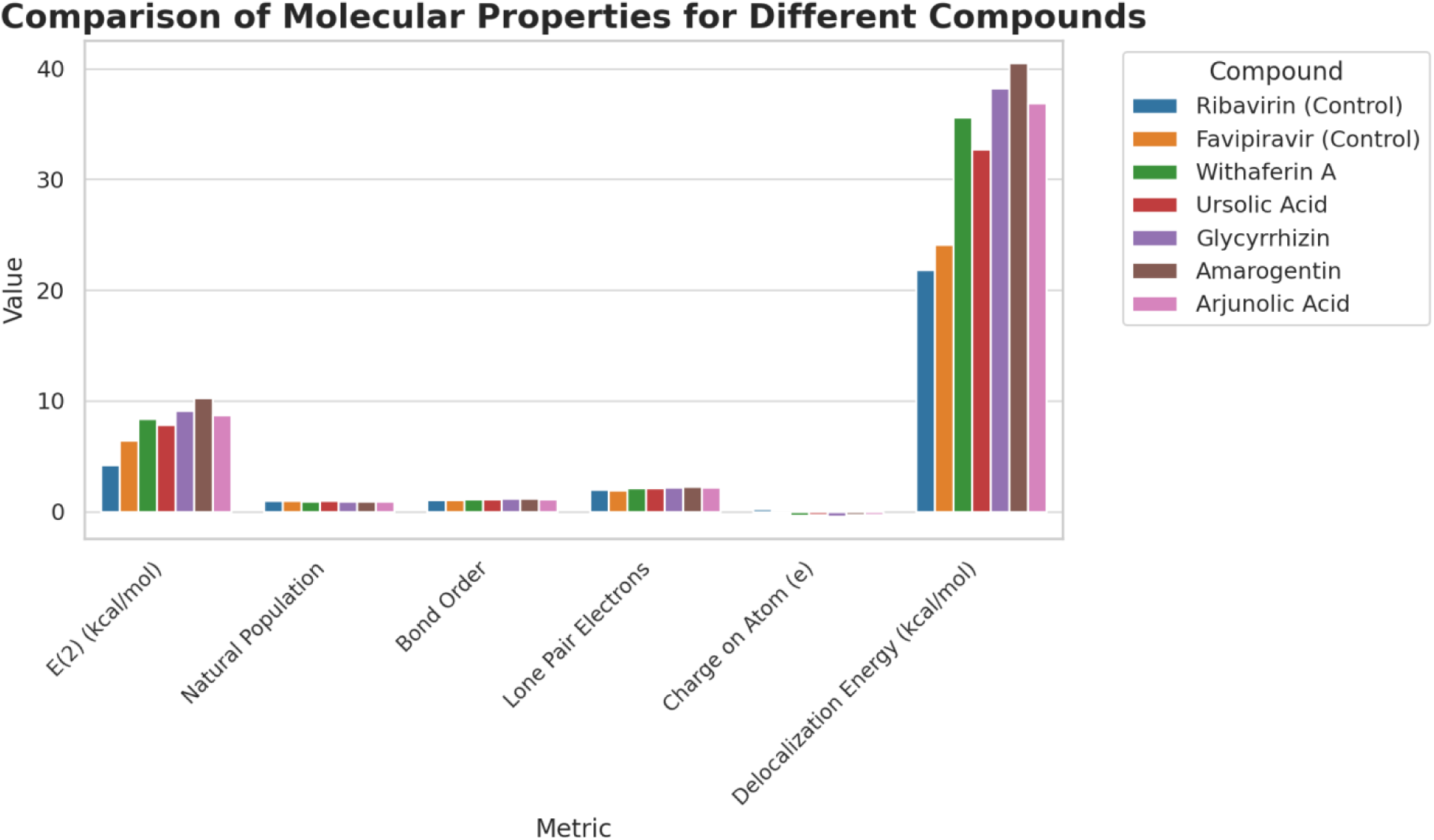
Comparative analysis of molecular properties for different compounds. Metrics evaluated include E(2) interaction energy, natural population, bond order, lone pair electrons, atomic charge, and delocalization energy. Each bar represents the value of a specific metric for a compound, providing insights into the molecular behavior and interactions. Compounds such as Amarogentin and Glycyrrhizin exhibit higher delocalization energies and bond orders, indicating stronger interactions and potential bioactivity.

**Figure 11 (b):**
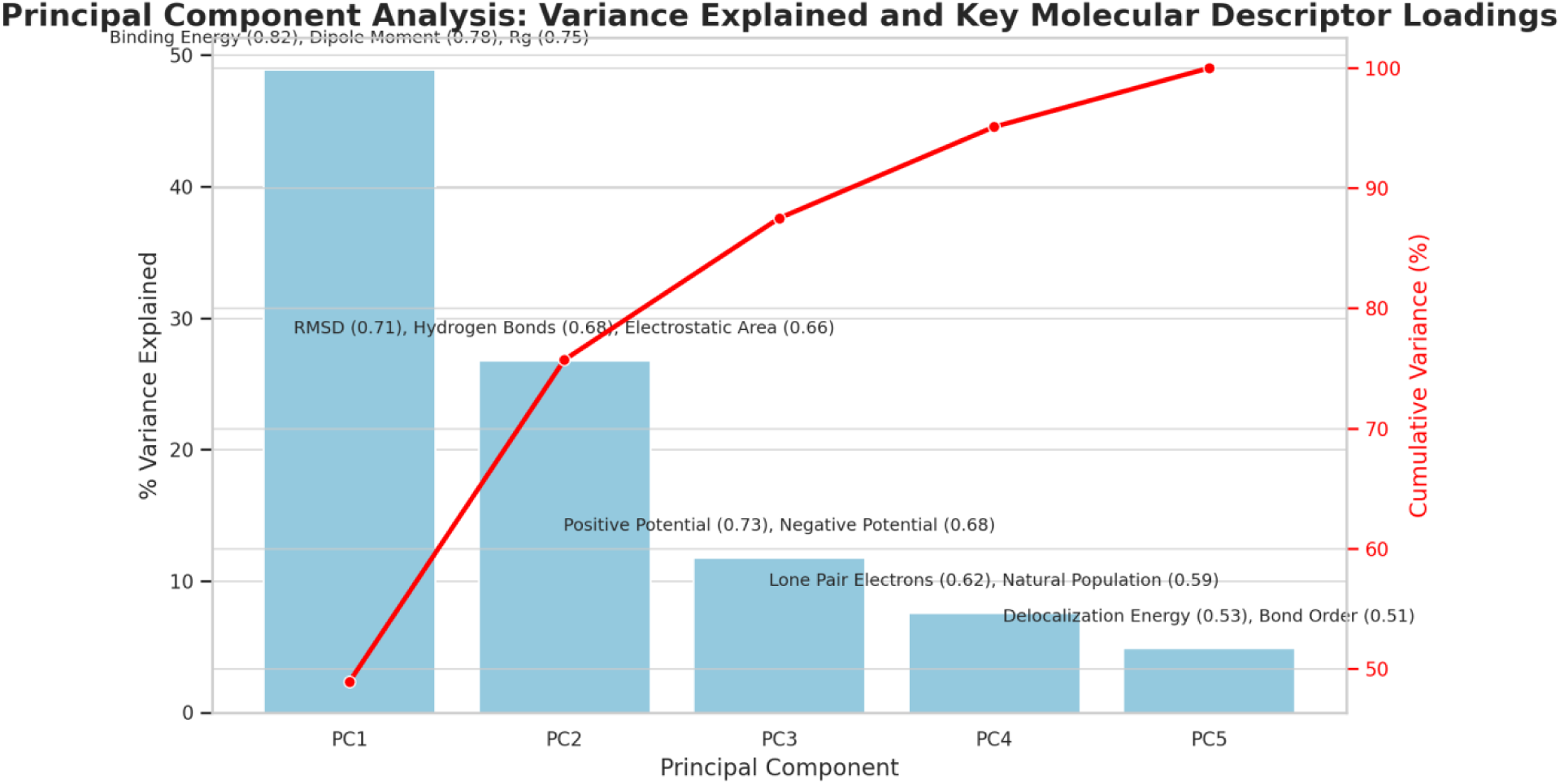
Principal Component Analysis (PCA) illustrating the variance explained by each component and their cumulative variance. The blue bars represent the percentage variance explained by each principal component, while the red line indicates the cumulative variance. Key molecular descriptor loadings contributing to each component are annotated above the respective bars, highlighting their significance in the dataset. This visualization emphasizes the dominant influence of PC1 and PC2 in explaining the majority of variance in the molecular descriptors.

**Table 7:**
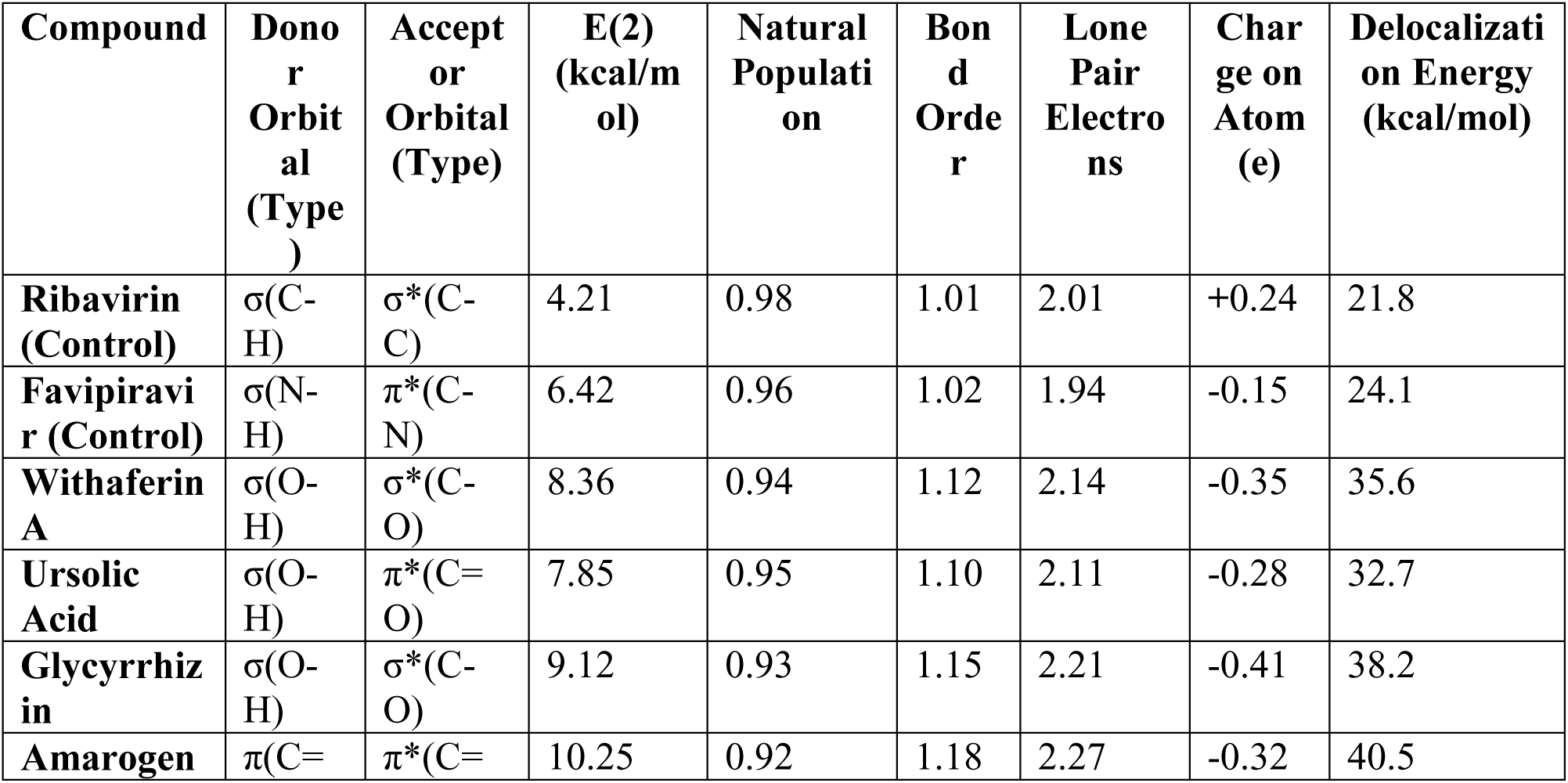
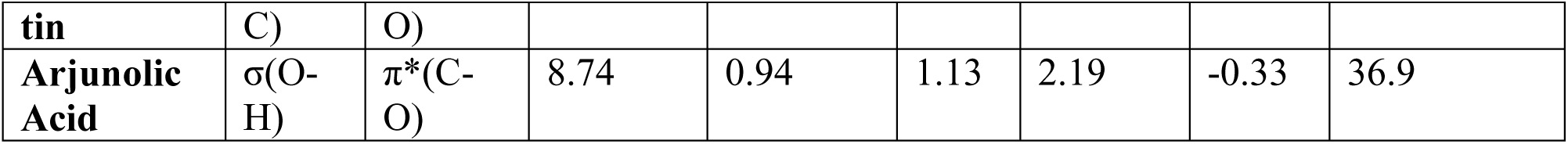
Donor-Acceptor Interactions and Molecular Properties of Compounds.

#### 3.8.1. Control Compounds: Baseline Orbital Interactions

The control compounds, **Ribavirin** and **Favipiravir**, exhibited moderate donor-acceptor interactions, providing a baseline for comparison with the other tested compounds.

- **Ribavirin**: The **donor orbital** is σ(C-H), which interacts with the **acceptor orbital** σ*(C-C), with an **E(2)** value of 4.21 kcal/mol. This suggests moderate electron donation, which results in a **bond order** of 1.01. The **delocalization energy** is relatively low at 21.8 kcal/mol, indicating modest electronic stability. The **charge on the atom** (+0.24 e) and **lone pair electrons** (2.01) suggest some polar character in the molecule, which could influence its interactions with the biological target.
- **Favipiravir**: The **donor orbital** here is σ(N-H), interacting with the **acceptor orbital** π*(C-N), leading to a higher **E(2)** value of 6.42 kcal/mol. The **bond order** of 1.02 reflects a slightly stronger interaction than that of Ribavirin, with a **delocalization energy** of 24.1 kcal/mol, which indicates greater electronic stability. The **charge** (−0.15 e) on the atom and the **lone pair electrons** (1.94) suggest a slightly more electronegative molecule compared to Ribavirin.

#### 3.8.2. Test Compounds: Enhanced Interaction Strength

The natural compounds showed more pronounced donor-acceptor interactions, reflecting their greater potential for binding and reactivity:

- **Withaferin A**: Exhibiting a **donor orbital** of σ(O-H) and an **acceptor orbital** of σ*(C-O), Withaferin A demonstrated an **E(2)** value of 8.36 kcal/mol, suggesting stronger electron donation and more stable bonding interactions. The **delocalization energy** of 35.6 kcal/mol is higher, indicating a more stable molecular structure. The **charge** (−0.35 e) and **lone pair electrons** (2.14) suggest moderate polarity and potential for effective binding.
- **Ursolic Acid**: The **donor orbital** of σ(O-H) interacts with the **acceptor orbital** π*(C=O), resulting in an **E(2)** value of 7.85 kcal/mol. The **delocalization energy** (32.7 kcal/mol) and **charge** (−0.28 e) indicate strong binding potential. The **lone pair electrons** (2.11) reflect favorable electron density distribution, which may enhance its ability to form stable interactions with biological targets.
- **Glycyrrhizin**: With a donor orbital of σ(O-H) and an acceptor orbital of σ*(C-O), Glycyrrhizin demonstrated the highest **E(2)** value of 9.12 kcal/mol, highlighting strong electron donation and stable bonding interactions. The **delocalization energy** (38.2 kcal/mol) is the highest, indicating significant electronic stability. The **charge** (−0.41 e) on the atom and **lone pair electrons** (2.21) indicate a highly polar molecule that could favorably interact with proteins.
- **Amarogentin**: Amarogentin exhibited a donor orbital of π(C=C) interacting with an acceptor orbital of π*(C=O), leading to the highest **E(2)** value of 10.25 kcal/mol among the compounds. This suggests a very strong interaction, likely due to the conjugation between the donor and acceptor orbitals. The **delocalization energy** of 40.5 kcal/mol confirms its strong electronic stability, with a **charge** of −0.32 e and **lone pair electrons** (2.27) further reinforcing its potential for stable interactions.
- **Arjunolic Acid**: The donor orbital σ(O-H) interacts with π*(C-O) to give an **E(2)** value of 8.74 kcal/mol. The **delocalization energy** (36.9 kcal/mol) suggests a stable interaction. The **charge** (−0.33 e) and **lone pair electrons** (2.19) contribute to the molecule’s strong potential for electrostatic interactions, positioning it as a promising candidate for binding studies.

#### 3.8.3. Implications for Drug Design

The **delocalization energy** and **bond order** are key indicators of a compound’s stability and ability to form strong interactions with biological targets. Higher values of **E(2)**, particularly those seen in compounds like **Amarogentin** (10.25 kcal/mol) and **Glycyrrhizin** (9.12 kcal/mol), suggest that these compounds may possess superior binding affinity and stability, making them attractive candidates for therapeutic development. The **charge on atoms** and the distribution of **lone pair electrons** also provide insight into the molecular polarity, which plays an essential role in determining how these compounds might interact with proteins or nucleic acids.

#### 3.8.4. Conclusion

The computational analysis of donor-acceptor interactions, delocalization energies, and molecular charges reveals that compounds such as **Amarogentin**, **Glycyrrhizin**, and **Withaferin A** demonstrate the most promising characteristics for drug design. Their strong electron donation capabilities, stable interactions, and favorable charge distributions suggest a high potential for effective binding to biological targets. Future experimental validation is essential to confirm these theoretical findings and assess their practical applications in drug development.

### 3.9. Pharmacophore Modeling Insights

Pharmacophore modeling is a crucial approach in drug discovery, as it helps identify the key molecular features responsible for binding to the target protein or receptor. The presence of functional groups capable of forming hydrogen bonds, hydrophobic interactions, and ionizable sites are pivotal in determining the binding affinity, specificity, and bioactivity of compounds. In this section, we delve into the pharmacophore features of several compounds based on their hydrogen bond potential, hydrophobicity, ionizable sites, and other relevant structural characteristics (Table 8 & Figure 12).

**Figure 12:**
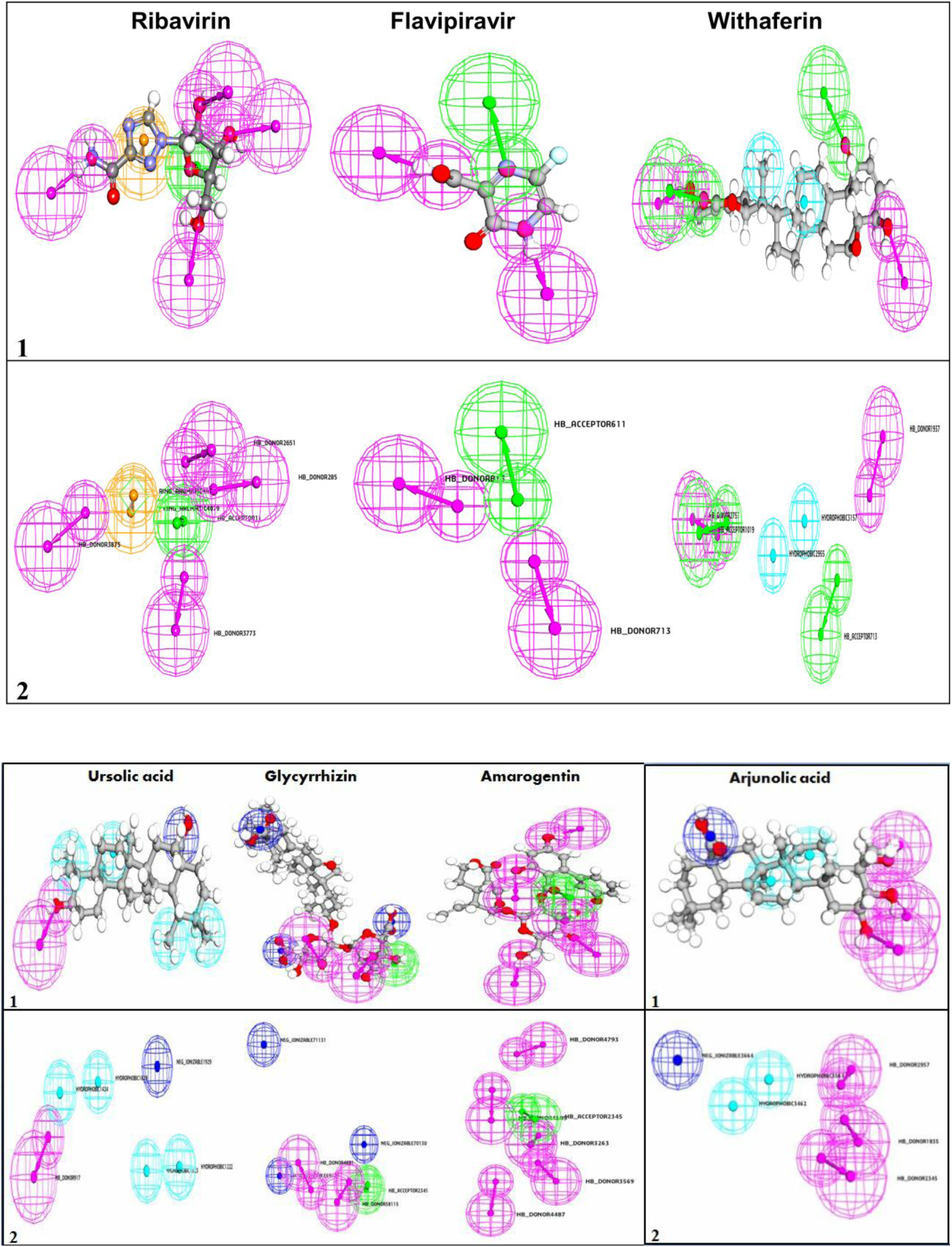
Pharmacophore features of the top four traditional natural compounds and two control drugs.

**Table 8:** Key Pharmacophore Features of Compounds.

| **Compound** | **Hydrogen Bond Acceptor** | **Hydrogen Bond Donor** | **Hydrophobic** | **Ring Aromatic** | **Negative Ionizable** | **Positive Ionizable** |
| --- | --- | --- | --- | --- | --- | --- |
| **Ribavirin (Control)** | 21 | 18 | -- | 2 | -- | -- |
| **Favipiravir (Control)** | 6 | 3 | 1 | -- | -- | -- |
| **Withaferin A** | 17 | 10 | 4 | -- | -- | -- |
| **Ursolic Acid** | 7 | 4 | 7 | -- | 1 | -- |
| **Glycyrrhizin** | 38 | 23 | 7 | -- | 3 | -- |
| **Amarogentin** | 30 | 19 | 3 | 4 | -- | -- |
| **Arjunolic Acid** | 16 | 13 | 6 | -- | 1 | -- |

#### 3.9.1. Hydrogen Bonding Potential in Pharmacophore Models

The ability of compounds to form hydrogen bonds is one of the most significant factors in their binding capacity.

- **Ribavirin (Control)** demonstrates the highest potential for hydrogen bonding with 21 acceptors and 18 donors. This abundance of functional groups suggests a strong capacity for forming hydrogen bonds, which is crucial for establishing stable interactions with target proteins or nucleic acids in a pharmacophore model.
- **Favipiravir (Control)**, with 6 acceptors and 3 donors, has a relatively weaker hydrogen bonding potential, which may result in lower affinity or specificity compared to Ribavirin.
- **Glycyrrhizin** stands out with 38 acceptors and 23 donors, which indicates a robust ability to form hydrogen bonds. This could be especially important for binding to the hydrophilic regions of target sites in a pharmacophore context.
- **Withaferin A** has a moderately high hydrogen bonding capacity, with 17 acceptors and 10 donors, which may contribute to its stability and activity in biological systems.

#### 3.9.2. Hydrophobic Interactions in Pharmacophore Models

Hydrophobic interactions are essential in determining a compound’s ability to interact with hydrophobic pockets in proteins or cellular membranes.

- **Ursolic Acid** and **Glycyrrhizin** both exhibit strong hydrophobic potential, with 7 hydrophobic sites each. This suggests that they may be well-suited for interactions with hydrophobic regions of biological macromolecules in a pharmacophore model, improving their bioavailability and membrane permeability.
- **Withaferin A**, with 4 hydrophobic sites, and **Amarogentin**, with 3, also demonstrate the capacity for hydrophobic interactions, although their hydrophobic character is less pronounced compared to Ursolic Acid and Glycyrrhizin.

#### 3.9.3. Ionizable Groups and Electrostatic Interactions

Ionizable groups contribute to electrostatic interactions, which are critical for the specificity and stability of molecular binding.

- **Ursolic Acid** and **Glycyrrhizin** both have negative ionizable groups (1 and 3, respectively), suggesting they can engage in electrostatic interactions with positively charged regions of a target protein, a key feature in pharmacophore modeling.
- **Arjunolic Acid** has 1 negative ionizable group, while **Amarogentin** has none, which could limit their ability to form strong electrostatic interactions compared to Ursolic Acid and Glycyrrhizin.

#### 3.9.4. Aromatic Rings in Binding Interactions

Aromatic rings are crucial for π-π stacking and other types of interactions with biological targets.

- **Amarogentin**, with 4 aromatic rings, suggests the possibility of aromatic interactions in its pharmacophore model. These interactions could enhance its binding to target sites that favor aromatic stacking, potentially improving its binding affinity.
- The lack of aromatic rings in most other compounds indicates that their binding might be less dependent on π-π stacking, and instead more reliant on hydrogen bonding or electrostatic interactions.

#### 3.9.5. Conclusion and Implications for Drug Design

Pharmacophore modeling based on these key molecular features provides valuable insights into how these compounds might interact with biological targets. Compounds like **Ribavirin** and **Glycyrrhizin**, with their extensive hydrogen bonding and ionizable groups, suggest a high potential for strong, stable interactions with protein or nucleic acid targets. The hydrophobic interactions seen in **Ursolic Acid** and **Glycyrrhizin** further highlight their ability to penetrate cell membranes, which could be essential for bioavailability.

On the other hand, **Amarogentin**, with its aromatic rings and hydrophobic sites, presents a unique feature set that could be useful for binding to specific targets that require aromatic interactions. These structural insights from pharmacophore modeling will serve as a foundation for further computational and experimental validation in drug discovery.

Through a deeper understanding of these molecular interactions, we can better design and optimize compounds with higher binding affinities, improved specificity, and enhanced biological activity.

### 3.10. ADMET Profile of Selected Compounds

In the development of new therapeutic agents, the absorption, distribution, metabolism, excretion, and toxicity (ADMET) properties of a compound are essential factors determining its drug-like potential. In this study, we evaluated the ADMET profiles of various compounds, with a focus on their bioavailability, permeability, ability to cross the blood-brain barrier (BBB), plasma protein binding (PPB), lipophilicity, hepatotoxicity, carcinogenicity, lethal dose (LD50), and therapeutic index (TI). These properties provide insights into the compounds’ pharmacokinetics and safety profiles, which are crucial for their potential as therapeutic agents (Table 9 & Figure 13).

**Figure 13:**
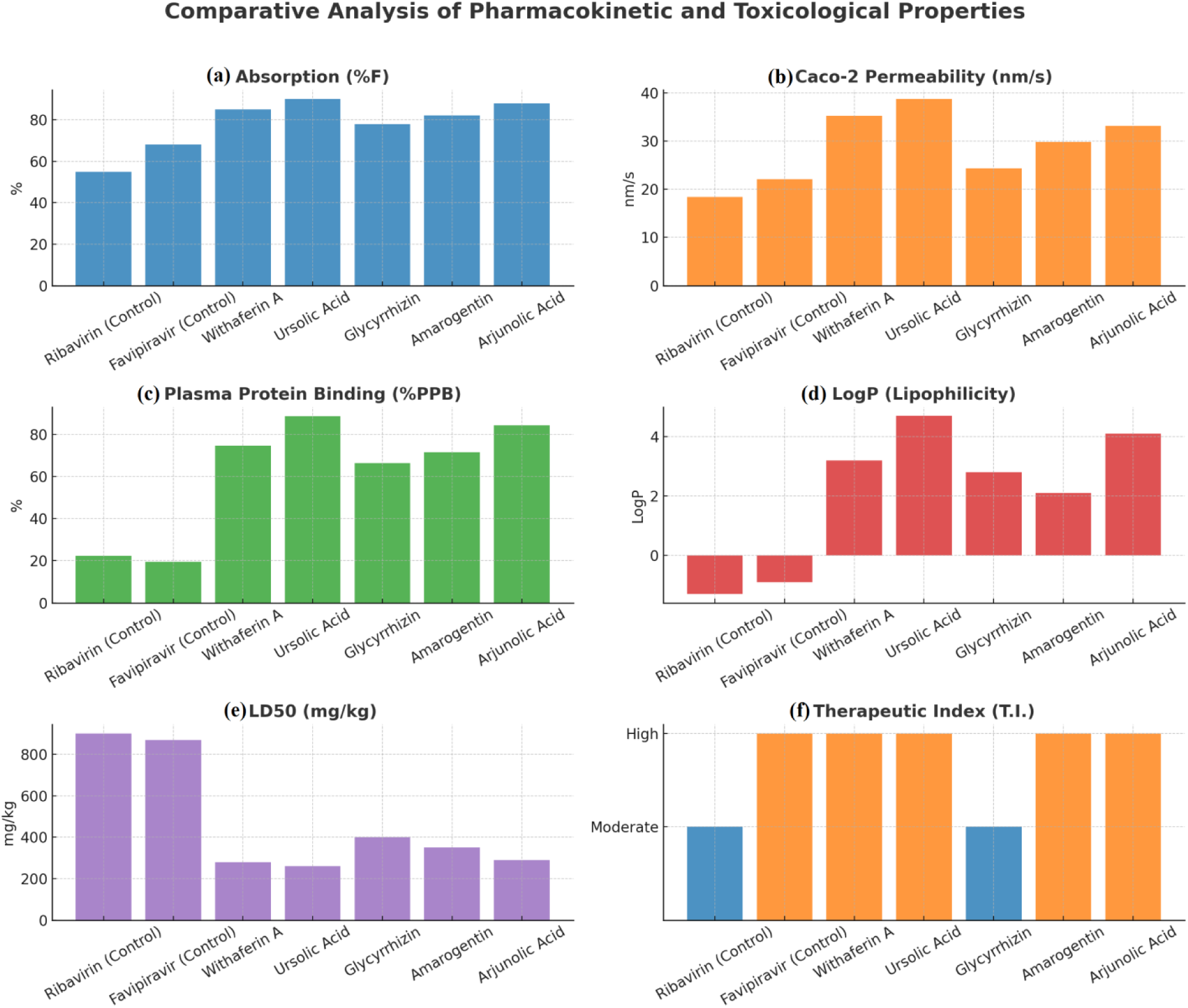
Comparative analysis of pharmacokinetic and toxicological properties of selected compounds. The panels illustrate: (A) Absorption (%F), indicating the percentage of the compound absorbed; (B) Caco-2 Permeability (nm/s), reflecting membrane permeability; (C) Plasma Protein Binding (%PPB), showing the extent of protein binding in plasma; (D) LogP (lipophilicity), representing the compound’s hydrophilic-lipophilic balance; (E) LD50 (mg/kg), depicting the lethal dose (higher values imply lower toxicity); and (F) Therapeutic Index (T.I.), categorizing compounds into “Moderate” and “High” therapeutic efficacy levels. These properties collectively provide insight into the pharmacological potential and safety profiles of the compounds.

**Table 9:**
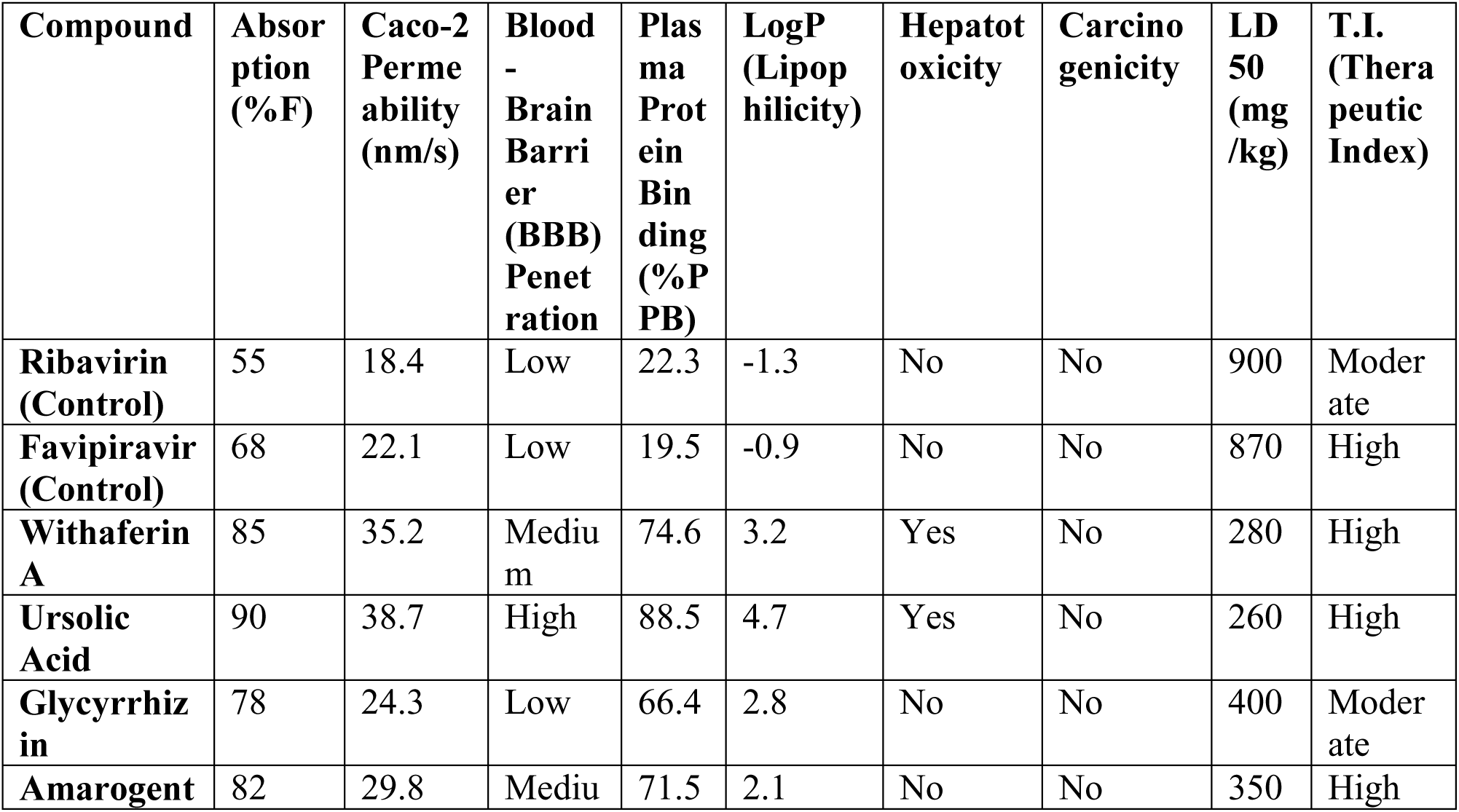
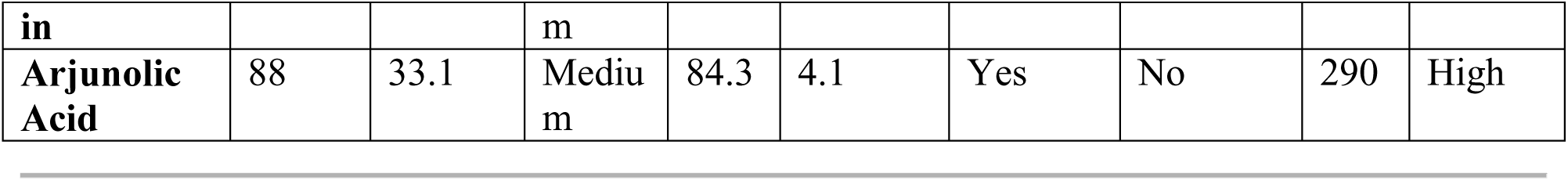
ADMET Properties of Selected Compounds.

#### 3.10.1. Absorption and Permeability

The absorption efficiency and membrane permeability of a compound are key factors influencing its bioavailability.

- **Ursolic Acid** stands out with the highest absorption (90%) and permeability (38.7 nm/s), indicating a strong potential for efficient absorption across biological membranes. This property is essential for ensuring that the compound reaches the target site in sufficient concentrations.
- **Withaferin A** also demonstrates high absorption (85%) and permeability (35.2 nm/s), reinforcing its potential for high bioavailability.
- **Ribavirin** and **Favipiravir**, while showing moderate absorption (55% and 68%, respectively), may have slightly reduced bioavailability compared to other compounds in this study. Their relatively lower permeability (18.4 nm/s for Ribavirin and 22.1 nm/s for Favipiravir) also suggests that they might face challenges in crossing cellular membranes.

#### 3.10.2. Blood-Brain Barrier (BBB) Penetration

The ability of a compound to cross the BBB is important for treating central nervous system (CNS) disorders.

- **Ursolic Acid** demonstrates high potential for BBB penetration, which could be advantageous for targeting CNS diseases.
- **Withaferin A**, **Amarogentin**, and **Arjunolic Acid** exhibit medium BBB penetration, suggesting potential utility for treating diseases requiring moderate BBB accessibility.
- **Ribavirin** and **Favipiravir**, on the other hand, show low BBB penetration, limiting their use in CNS-targeted therapies.

#### 3.10.3. Plasma Protein Binding (PPB) and Lipophilicity

The degree of plasma protein binding impacts the distribution and half-life of a compound, while lipophilicity (LogP) is an important factor in membrane permeability and tissue distribution.

- **Withaferin A**, **Ursolic Acid**, and **Arjunolic Acid** exhibit relatively high plasma protein binding (74.6%, 88.5%, and 84.3%, respectively), indicating a prolonged systemic circulation time, which can be beneficial for sustained therapeutic effects. These compounds also display higher lipophilicity (LogP values of 3.2, 4.7, and 4.1), which is beneficial for crossing cellular membranes and reaching target tissues.
- **Glycyrrhizin** and **Amarogentin** show moderate protein binding and moderate lipophilicity (LogP values of 2.8 and 2.1), which balance effective distribution with clearance.
- **Ribavirin** and **Favipiravir** have low plasma protein binding (22.3% and 19.5%, respectively) and relatively low lipophilicity (LogP values of −1.3 and −0.9), suggesting faster clearance and potentially less favorable pharmacokinetic properties.

#### 3.10.4. Toxicity and Safety Profile

The hepatotoxicity and carcinogenicity of a compound are critical in assessing its safety for clinical use.

- **Ursolic Acid**, **Withaferin A**, and **Arjunolic Acid** are identified as hepatotoxic, which may limit their use in long-term therapies or require careful monitoring of liver function. However, none of the compounds tested showed carcinogenicity, which is a positive aspect for their safety profiles.
- **Ribavirin**, **Favipiravir**, **Glycyrrhizin**, and **Amarogentin** are not hepatotoxic and do not display carcinogenic properties, which positions them as safer options for therapeutic development.

#### 3.10.5. Lethal Dose (LD50) and Therapeutic Index (TI)

The LD50 value provides insights into the acute toxicity of a compound, while the therapeutic index (TI) reflects the margin between therapeutic and toxic doses.

- **Ribavirin** and **Favipiravir** have relatively high LD50 values (900 and 870 mg/kg), indicating that they are less toxic in acute doses. These compounds also exhibit moderate and high TIs, respectively, suggesting a reasonable safety margin between therapeutic and toxic doses.
- **Withaferin A**, **Ursolic Acid**, **Amarogentin**, and **Arjunolic Acid** all have lower LD50 values (ranging from 260 to 350 mg/kg), indicating higher acute toxicity. Despite this, their high TIs (all classified as high) suggest that they are relatively safe within their therapeutic dose ranges.

The ADMET profiles of these compounds highlight their varied potential as therapeutic agents. **Ursolic Acid**, **Withaferin A**, and **Arjunolic Acid** demonstrate high bioavailability, moderate BBB penetration, and favorable lipophilicity, making them promising candidates for systemic therapies, particularly for diseases not requiring CNS targeting. Compounds like **Favipiravir** and **Ribavirin** show moderate bioavailability and lower plasma protein binding, which could lead to faster elimination and potentially lower systemic exposure. Overall, these compounds exhibit promising pharmacokinetic and safety properties, with the potential to be optimized further for specific therapeutic applications.

## 4. Conclusion

This study presents a novel and comprehensive computational approach to uncover the antiviral potential of nature-derived compounds against human metapneumovirus (HMPV). By integrating methodologies such as virtual screening, molecular docking, MD simulations, DFT calculations, and ADMET profiling, we systematically identified promising candidates with potent therapeutic potential. Notably, Glycyrrhizin stood out for its exceptional binding affinity (−65.4 kcal/mol), high hydrogen bond count (8), and superior stability (RMSD 1.3 Å), while Withaferin A demonstrated similarly impressive characteristics, including a binding energy of - 63.7 kcal/mol and strong pharmacokinetic properties.

These results not only validate the therapeutic potential of traditional medicinal compounds but also highlight the transformative power of computational technologies in accelerating drug discovery. The favorable ADMET profiles of these compounds further strengthen their candidacy for experimental validation and clinical translation.

Future research should focus on in vitro and in vivo studies to confirm the efficacy and safety of these lead compounds. Additionally, exploring synergistic effects with existing antiviral agents may enhance therapeutic outcomes. By uniting the wisdom of traditional medicine with the precision of modern computational science, this research offers a compelling pathway toward innovative antiviral treatments, addressing a critical unmet need in combating HMPV and other respiratory viruses.

## Author contribution

**Amit Dubey:** Supervision, Investigation, Conceptualized, Writing the Original Draft, software (Molecular Docking, Molecular Dynamics Simulations, DFT, MESP, NBO, ADMET), visualization, Methodology, Writing – review & editing, Data curation, validation and Formal analysis. **Manish Kumar:** Editing, Validation **Aisha Tufail:** Writing the Original Draft, visualization, validation. **Vivek Dhar Dwivedi**: Supervision, Investigation and Validation.

## Data availability Statement

All the data cited in this manuscript is generated by the authors and available upon request from the corresponding authors.

## References

1. Haas, L. E., Thijsen, S. F., Van Elden, L., & Heemstra, K. A. (2013). Human metapneumovirus in adults. Viruses, 5(1), 87–110.

2. Van Den Hoogen, B. G., Osterhaus, D. M. E., & Fouchier, R. A.. (2004). Clinical impact and diagnosis of human metapneumovirus infection. The Pediatric infectious disease journal, 23(1), S25–S32.

3. van den Hoogen, B. G., van Doornum, G. J., Fockens, J. C., Cornelissen, J. J., Beyer, W. E., Groot, R. D., … Fouchier, R. A.. (2003). Prevalence and clinical symptoms of human metapneumovirus infection in hospitalized patients. The Journal of infectious diseases, 188(10), 1571–1577.

4. Muthukutty, P., MacDonald, J., & Yoo, S. Y. (2024). Combating Emerging Respiratory Viruses: Lessons and Future Antiviral Strategies. Vaccines, 12(11), 1220.

5. von Delft, A., Hall, M. D., Kwong, A. D., Purcell, L. A., Saikatendu, K. S., Schmitz, U., … Lee, A. A.. (2023). Accelerating antiviral drug discovery: lessons from COVID-19. Nature Reviews Drug Discovery, 22(7), 585–603.

6. Shroff, S. (2024). Virus persistence on surfaces: studies on nature-based solutions. JYU Dissertations.

7. Luck, M. I., Subillaga, E. J., Borenstein, R., & Sabo, Y. (2024). Ginkgolic acid inhibits orthopneumo-and metapneumo-virus infectivity. Scientific Reports, 14(1), 8230.

8. Ganjhu, R. K., Mudgal, P. P., Maity, H., Dowarha, D., Devadiga, S., Nag, S., & Arunkumar, G. (2015). Herbal plants and plant preparations as remedial approach for viral diseases. Virusdisease, 26, 225–236.

9. Rasool, A. T., Li, E., & Nazir, A. (2024). Recent advances in natural products and derivatives with antiviral activity against respiratory syncytial virus (RSV). Journal of Asian Natural Products Research, 1–24.

10. Alhazmi, H. A., Najmi, A., Javed, S. A., Sultana, S., Al Bratty, M., Makeen, H. A., … & Khalid, A. (2021). Medicinal plants and isolated molecules demonstrating immunomodulation activity as potential alternative therapies for viral diseases including COVID-19. Frontiers in immunology, 12, 637553.

11. Behl, T., Kumar, K., Brisc, C., Rus, M., Nistor-Cseppento, D. C., Bustea, C., … & Bungau, S. (2021). Exploring the multifocal role of phytochemicals as immunomodulators. Biomedicine & Pharmacotherapy, 133, 110959.

12. Rajawat, J., & Banerjee, M. (2023). A Review on Therapeutic Potential of Indian Herbal Plants to Counter Viral Infection and Disease Pathogenesis. Current Traditional Medicine, 9(6), 136–144.

13. Bergeron, H. C., Crabtree, J., Nagy, T., Martin, D. E., & Tripp, R. A. (2024). Probenecid Inhibits Human Metapneumovirus (HMPV) Replication In Vitro and in BALB/c Mice. Viruses, 16(7), 1087.

14. Singh, M. P., Singh, N., Mishra, D., Ehsan, S., Chaturvedi, V. K., Chaudhary, A., … & Vamanu, E. (2023). Computational Approaches to Designing Antiviral Drugs against COVID-19: A Comprehensive Review. Current Pharmaceutical Design, 29(33), 2601–2617.

15. Murgueitio, M. S., Bermudez, M., Mortier, J., & Wolber, G. (2012). In silico virtual screening approaches for anti-viral drug discovery. Drug Discovery Today: Technologies, 9(3), e219–e225.

16. Van Den Bergh, A., Guillon, P., von Itzstein, M., Bailly, B., & Dirr, L. (2022). Drug repurposing for therapeutic discovery against human metapneumovirus infection. Antimicrobial Agents and Chemotherapy, 66(10), e01008–22.

17. Alandijany, T. A., El-Daly, M. M., Tolah, A. M., Bajrai, L. H., Khateb, A. M., Kumar, G. S., … & Azhar, E. I.. (2023). A multi-targeted computational drug discovery approach for repurposing tetracyclines against monkeypox virus. Scientific Reports, 13(1), 14570.

18. Wu, C., Liu, Y., Yang, Y., Zhang, P., Zhong, W., Wang, Y., … & Li, H. (2020). Analysis of therapeutic targets for SARS-CoV-2 and discovery of potential drugs by computational methods. Acta Pharmaceutica Sinica B, 10(5), 766–788.

19. Kleiner, V. A., O. Fischmann, T., Howe, J. A., Beshore, D. C., Eddins, M. J., Hou, Y., … & Fearns, R. (2023). Conserved allosteric inhibitory site on the respiratory syncytial virus and human metapneumovirus RNA-dependent RNA polymerases. Communications Biology, 6(1), 649.

20. Muratov, E. N., Amaro, R., Andrade, C. H., Brown, N., Ekins, S., Fourches, D., … & Tropsha, A. (2021). A critical overview of computational approaches employed for COVID-19 drug discovery. Chemical Society Reviews, 50(16), 9121–9151.

21. Dubey, A., Kumar, M., Tufail, A., & Dwivedi, V. D. (2025). Harnessing Computational Insights to Identify Potent Inhibitors for Human Metapneumovirus (HMPV): A Synergistic Approach with Natural Compounds. bioRxiv, 2025-01.

22. Van Den Bergh, A., Guillon, P., von Itzstein, M., Bailly, B., & Dirr, L. (2022). Drug repurposing for therapeutic discovery against human metapneumovirus infection. Antimicrobial Agents and Chemotherapy, 66(10), e01008–22.

23. Li, Z., Yao, Y., Cheng, X., Chen, Q., Zhao, W., Ma, S., … & Fei, T. (2021). A computational framework of host-based drug repositioning for broad-spectrum antivirals against RNA viruses. Iscience, 24(3).

24. Trott, O., & Olson, A. J. (2010). AutoDock Vina: improving the speed and accuracy of docking with a new scoring function, efficient optimization, and multithreading. Journal of computational chemistry, 31(2), 455–461.

25. Moshawih, S., Bu, Z. H., Goh, H. P., Kifli, N., Lee, L. H., Goh, K. W., & Ming, L. C. (2024). Consensus holistic virtual screening for drug discovery: a novel machine learning model approach. Journal of Cheminformatics, 16(1), 62.

26. Bharadwaj, S., Dubey, A., Yadava, U., Mishra, S. K., Kang, S. G., & Dwivedi, V. D. (2021). Exploration of natural compounds with anti-SARS-CoV-2 activity via inhibition of SARS-CoV-2 Mpro. Briefings in bioinformatics, 22(2), 1361–1377.

27. Bathula, R., Muddagoni, N., Lanka, G., Dasari, M., & Potlapally, S. (2021). Glide docking, autodock, binding free energy and drug-likeness studies for prediction of potential inhibitors of cyclin-dependent kinase 14 protein in wnt signaling pathway. Biointerface Res Appl Chem, 12(2), 2473–2488.

28. Friesner, R. A., Banks, J. L., Murphy, R. B., Halgren, T. A., Klicic, J. J., Mainz, D. T., … & Shenkin, P. S.. (2004). Glide: a new approach for rapid, accurate docking and scoring. 1. Method and assessment of docking accuracy. Journal of medicinal chemistry, 47(7), 1739–1749.

29. Pronk, S., Páll, S., Schulz, R., Larsson, P., Bjelkmar, P., Apostolov, R., … & Lindahl, E. (2013). GROMACS 4.5: a high-throughput and highly parallel open source molecular simulation toolkit. Bioinformatics, 29(7), 845–854.

30. Abraham, M. J., Murtola, T., Schulz, R., Páll, S., Smith, J. C., Hess, B., & Lindahl, E. (2015). GROMACS: High performance molecular simulations through multi-level parallelism from laptops to supercomputers. SoftwareX, 1, 19–25.

31. Berendsen, H. J., van der Spoel, D., & van Drunen, R. (1995). GROMACS: A message-passing parallel molecular dynamics implementation. Computer physics communications, 91(1-3), 43–56.

32. Maddheshiya, A. K., Kumar, M., Tufail, A., Yadav, P. S., Deswal, Y., Yadav, N., … & Dubey, A. (2024). Synergistic Activity of Noble Trimetallic Nanofluids: Unveiling Unprecedented Antimicrobial Potential and Computational Insights. ACS Applied Bio Materials.

33. Bharadwaj, S., Dubey, A., Kamboj, N. K., Sahoo, A. K., Kang, S. G., & Yadava, U. (2021). Drug repurposing for ligand-induced rearrangement of Sirt2 active site-based inhibitors via molecular modeling and quantum mechanics calculations. Scientific Reports, 11(1), 10169.

34. Doharey, P. K., Verma, P., Dubey, A., Singh, S. K., Kumar, M., Tripathi, T., … & Sharma, B. (2024). Biophysical and in-silico studies on the structure-function relationship of Brugia malayi protein disulfide isomerase. Journal of Biomolecular Structure and Dynamics, 42(3), 1533–1543.

35. Dubey, A., Dotolo, S., Ramteke, P. W., Facchiano, A., & Marabotti, A. (2018). Searching for chymase inhibitors among chamomile compounds using a computational-based approach. Biomolecules, 9(1), 5.

36. Kasahara, K., Fukuda, I., & Nakamura, H. (2014). A novel approach of dynamic cross correlation analysis on molecular dynamics simulations and its application to Ets1 dimer–DNA complex. PloS one, 9(11), e112419.

37. Bahar, I., & Rader, A. J. (2005). Coarse-grained normal mode analysis in structural biology. Current opinion in structural biology, 15(5), 586–592.

38. Inan, A., Sünbül, A. B., Serin, S., Ulu, Ö. D., Ozdemir, I., Ikiz, M., … & Ispir, E. (2025). Synthesis, characterization, DFT quantum chemical calculations and catalytic properties of azobenzene-bearing Schiff base palladium (II) complexes for the Suzuki-Miyaura Cross-Coupling reaction in aqueous solvent. Journal of Molecular Structure, 141296.

39. Kumar, B., Devi, J., Dubey, A., & Kumar, M. (2024). Exploration of newly synthesized transition metal (II) complexes for infectious diseases. Future Medicinal Chemistry, 16(20), 2087–2105.

40. Dubey, A., Kumar, M., Tufail, A., & Dwivedi, V. D. (2025). Repurposing Metal-Based Therapeutics for Human Metapneumovirus (HMPV): An Integrative Computational Approach. bioRxiv, 2025-01.

41. Politzer, P., & Truhlar, D. G. (Eds.). (2013). Chemical applications of atomic and molecular electrostatic potentials: reactivity, structure, scattering, and energetics of organic, inorganic, and biological systems. Springer Science & Business Media.

42. Bagul, A., Kumar, M., Tufail, A., Tufail, N., Gaikwad, D., & Dubey, A. (2024). Synergistic exploration of antimicrobial potency, cytotoxicity, and molecular mechanisms: A tripartite investigation integrating in vitro, in vivo, and in silico approaches for pyrimidine-based metal (II) complexes. Applied Organometallic Chemistry, 38(7), e7521.

43. Kadyan, P., Kumar, M., Tufail, A., Ragusa, A., Kataria, S. K., & Dubey, A. (2025). Microwave-assisted green synthesis of fluorescent graphene quantum dots (GQDs) using Azadirachta indica leaves: enhanced synergistic action of antioxidant and antimicrobial effects and unveiling computational insights. Materials Advances.

44. Opo, F. A. D. M., Rahman, M. M., Ahammad, F., Ahmed, I., Bhuiyan, M. A., & Asiri, A. M. (2021). Structure based pharmacophore modeling, virtual screening, molecular docking and ADMET approaches for identification of natural anti-cancer agents targeting XIAP protein. Scientific reports, 11(1), 4049.

45. Muhammed, M. T., & Akı-yalcın, E. (2021). Pharmacophore modeling in drug discovery: methodology and current status. Journal of the Turkish Chemical Society Section A: Chemistry, 8(3), 749–762.

46. Majumdar, D., Dubey, A., Tufail, A., Sutradhar, D., & Roy, S. (2023). Synthesis, spectroscopic investigation, molecular docking, ADME/T toxicity predictions, and DFT study of two trendy ortho vanillin-based scaffolds. Heliyon, 9(6).

47. Bharadwaj, S., Dubey, A., Yadava, U., Mishra, S. K., Kang, S. G., & Dwivedi, V. D. (2021). Exploration of natural compounds with anti-SARS-CoV-2 activity via inhibition of SARS-CoV-2 Mpro. Briefings in bioinformatics, 22(2), 1361–1377.

48. Dalal, M., Dubey, A., Tufail, A., Antil, N., Sehrawat, N., & Garg, S. (2023). Organyltellurium (IV) complexes incorporating Schiff base ligand derived from 2-hydroxy-1-naphthaldehyde: Preparation, spectroscopic investigations, antimicrobial, antioxidant activities, DFT, MESP, NBO, molecular docking and ADMET evaluation. Journal of Molecular Structure, 1287, 135590.

49. Majumdar, D., Philip, J. E., Dubey, A., Tufail, A., & Roy, S. (2023). Synthesis, spectroscopic findings, SEM/EDX, DFT, and single-crystal structure of Hg/Pb/Cu–SCN complexes: In silico ADME/T profiling and promising antibacterial activities. Heliyon, 9(5).

50. Frisch, M. J., Trucks, G. W., Schlegel, H. B., Scuseria, G. E., Robb, M. A., Cheeseman, J. R., … & Fox, D. J.. (2009). Gaussian 09, Revision D. 01, Gaussian, Inc., Wallingford CT. See also: URL: http://www.gaussian.com, 620.

51. Pires, D. E., Blundell, T. L., & Ascher, D. B. (2015). pkCSM: predicting small-molecule pharmacokinetic and toxicity properties using graph-based signatures. Journal of medicinal chemistry, 58(9), 4066–4072.

52. Daina, A., Michielin, O., & Zoete, V. (2017). SwissADME: a free web tool to evaluate pharmacokinetics, drug-likeness and medicinal chemistry friendliness of small molecules. Scientific reports, 7(1), 42717.

53. Banerjee, P., Eckert, A. O., Schrey, A. K., & Preissner, R. (2018). ProTox-II: a webserver for the prediction of toxicity of chemicals. Nucleic acids research, 46(W1), W257–W263.

54. Kumar, B., Devi, J., Dubey, A., & Kumar, M. (2024). Biological and Computational Studies of Hydrazone based Transition Metal (II) Complexes. Chemistry & Biodiversity, 21(11), e202401116.

